# A high-fat hypertensive diet induces a coordinated perturbation signature across cell types in thoracic perivascular adipose tissue

**DOI:** 10.1101/2025.02.13.636878

**Authors:** Leah Terrian, Janice M. Thompson, Derek E. Bowman, Vishal Panda, G. Andres Contreras, Cheryl E. Rockwell, Lisa Sather, Gregory D. Fink, D. Adam Lauver, Rance Nault, Stephanie W. Watts, Sudin Bhattacharya

**Affiliations:** Department of Biomedical Engineering, Michigan State University, East Lansing, MI, USA; Department of Pharmacology and Toxicology, Michigan State University, East Lansing, MI, USA; Institute for Quantitative Health Science and Engineering, Michigan State University, East Lansing, MI, USA; College of Osteopathic Medicine, Michigan State University, East Lansing, MI, USA; Department of Large Animal Clinical Sciences, Michigan State University, East Lansing, MI, USA; Institute for Integrative Toxicology, Michigan State University, East Lansing, MI, USA

## Abstract

Perivascular adipose tissue (PVAT), an intriguing layer of fat surrounding blood vessels, regulates vascular tone and mediates vascular dysfunction through mechanisms that are not well understood. Here we show with single nucleus RNA-sequencing of thoracic aortic PVAT from Dahl SS rats that a high-fat (HF) hypertensive diet induces coordinated changes in gene expression across the diverse cell types within PVAT. HF diet produced sex-specific alterations in cell-type proportions and genes related to remodeling of extracellular matrix dynamics and vascular integrity and stiffness, as well as changes in cell-cell communication pathways involved in angiogenesis, vascular remodeling, and mechanotransduction. Gene regulatory network analysis with virtual transcription factor knockout in adipocytes identified specific nuclear receptors that could be targeted for suppression or potential reversal of HF diet-induced changes. Interestingly, generative deep learning models were able to predict cross-cell-type perturbations in gene expression, indicating a hypertensive disease signature that characterizes HF-diet-induced perturbations in PVAT.

## INTRODUCTION

Cardiovascular diseases (CVD) were the cause of more than 20.5 million deaths globally in 2021 alone^1^. Hypertension is the most prevalent CVD with over 1 billion people affected and is considered the entree to other CVDs like heart failure, coronary artery disease, stroke, and peripheral artery disease^2^. Hypertension is especially problematic in obese and overweight subjects where the development of high blood pressure has been directly linked to the expansion of perivascular adipose tissue (PVAT)^3^. The PVAT, or tunica adiposa, is the outermost layer of tissue surrounding most blood vessels. Long considered to be a passive tissue involved in lipid storage and mechanical support, PVAT is now understood to be an active contributor to vascular function, playing important roles both in physiological homeostasis and its perturbation in metabolic and cardiovascular disease^4–9^. Adipocytes, the predominant cell type in PVAT, secrete various molecules including adipokines and cytokines, thereby modulating the functioning of smooth muscle and endothelial cells through paracrine signaling^5^. Additionally, *ex vivo* mechanical studies have shown that adipocytes play a key role in regulation of aortic stiffness^10^.

However, PVAT remains an enigmatic tissue whose precise mechanistic role in vascular dysfunction is not well understood. In healthy tissue, PVAT exerts an anticontractile effect on blood vessels, thereby helping maintain vascular tone and blood pressure homeostasis. However, this benefit of PVAT is lost in many cardiovascular diseases, including hypertension^4^, where changes in the physiological function of PVAT have been observed in arteries from both rodents and humans^11–13^. We focus here on possible alterations in PVAT phenotype in response to a high fat (HF) diet that elevates blood pressure in a rat model. The specific changes in gene expression and cell-cell communication induced by hypertension from excessive adiposity across the multiple cell types present in PVAT remain uncharacterized, hampering our overall understanding of the role of this tissue in cardiovascular disease. We address this knowledge gap by exploring several mechanistic questions. Specifically, how do the gene expression patterns in the diverse cell types in PVAT change when challenged with HF diet, and are these changes coordinated across cell types or unrelated? How do crucial patterns of cell-cell communication adapt in response to an HF diet? What gene regulatory networks mediate these changes? Are the observed changes sex-specific, and importantly, are they potentially reversible?

We examined these questions with novel single-nucleus RNA sequencing (snRNA-seq) analysis of thoracic aortic PVAT (taPVAT) from Dahl salt sensitive (Dahl SS) rats fed a HF diet. This rat model, developed by us and others^14^, reproducibly develops elevated blood pressure and increased visceral adiposity when fed a HF diet (60% kcal fat) from weaning, compared to age-matched rats fed a control diet (10% kcal fat). Moreover, the HF diet causes hypertension in both male and female Dahl SS rats. It is thus a good model for hypertension in humans, where up to 70% of individuals with hypertension are obese, defined as a Body Mass Index of 30 or above^15^.

We characterized time- and sex-specific alterations induced by HF diet in the proportions of different constituent cell types in PVAT: adipocytes, endothelial cells, fibroblasts, mesothelial cells, immune cells, pericytes, smooth muscle cells, and neuronal cells. Additionally, individual cell types showed characteristic changes in gene expression patterns induced by diet, in a manner dependent on both sex and duration on diet. Analyzing cell-cell communication among cell types in PVAT, we found broad similarities in inferred signaling pathways across sex and diet at the level of cell types in aggregate. However, there was considerable heterogeneity in signaling mechanisms both within and between cell types at the level of individual cells. Gene regulatory network analysis with CellOracle^16^ identified key transcriptional drivers of changes in adipocyte gene expression and revealed potential molecular targets to suppress or possibly reverse diet-dependent alterations in gene expression.

Finally, we show that a generative deep learning model^17^ was able to predict HF diet-induced changes in gene expression across cell types, essentially showing that there is a specific, quantifiable “perturbation signature” for diet-induced hypertension which is a function of cell type and basal gene expression. An overview of our workflow is shown in Figure 1.

**Figure 1.**
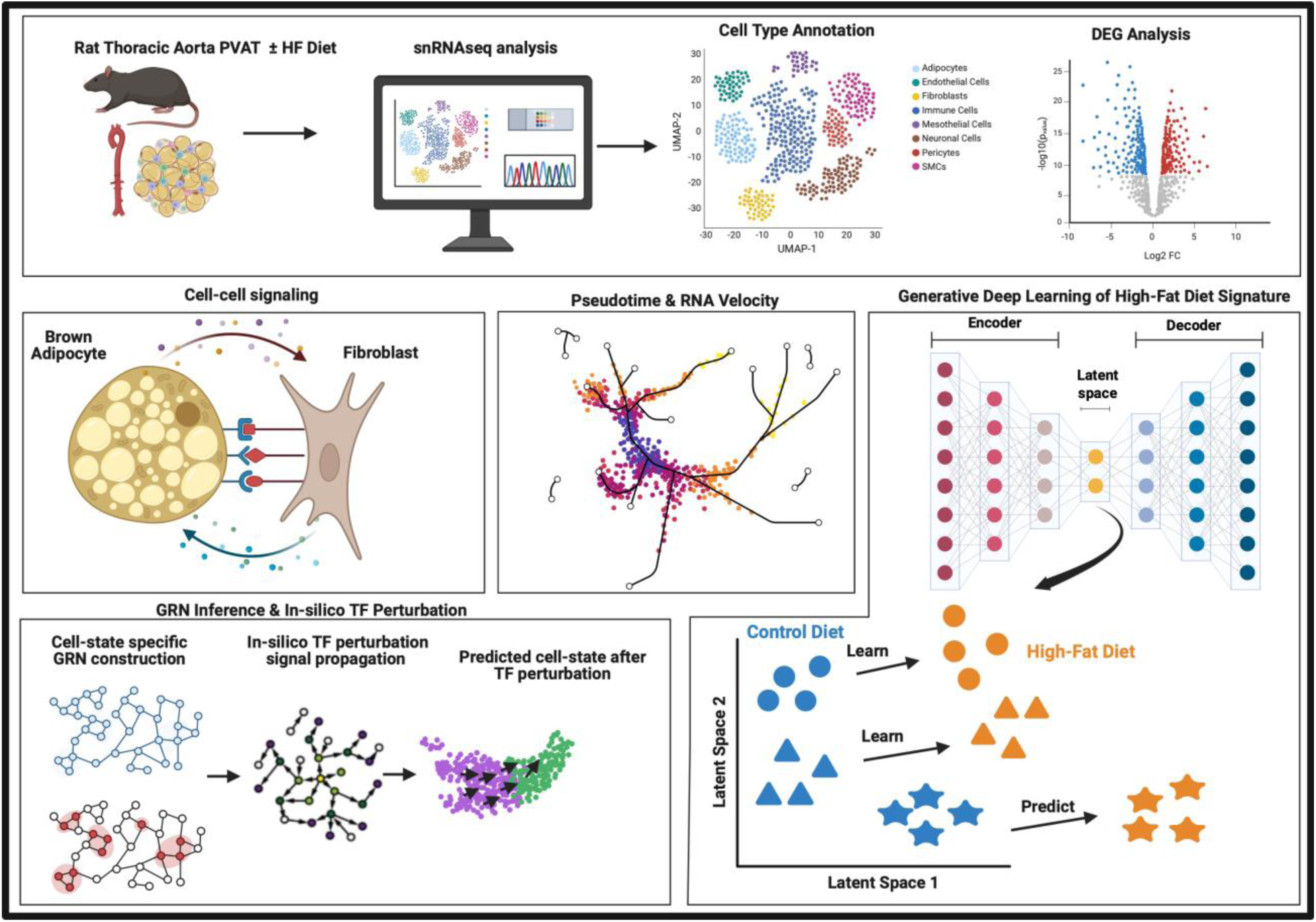
Overview of workflow and predictive modeling framework. Thoracic aorta perivascular adipose tissue (taPVAT) is collected from Dahl SS rats under control and high-fat (HF) diet conditions and profiled with snRNAseq, followed by clustering and annotation into major cell types. We then characterize condition-stratified differential gene expression induced by HF diet across cell types present in taPVAT. Next we infer putative cell-cell communication patterns based on ligand-receptor interactions between key cell types and identify directional shifts in cell state with pseudotime and RNA velocity analysis. In addition, we infer cell-state–specific gene regulatory networks (GRNs) using CellOracle^16^ and quantitatively predict cell state shifts with *in-silico* transcription factor perturbations. Finally, we use our previously developed deep generative learning model for single-cell perturbation prediction^17^ to identify an HF-diet-associated perturbation signature that could be quantitatively extrapolated to predict changes in gene expression across cell types. Created in BioRender. Bowman, D. (2026) https://BioRender.com/lxqxr83

## RESULTS

### Both male and female Dahl SS rats exhibit increased blood pressure when fed a high fat diet

We first evaluated the difference in body weight and mean arterial pressure (MAP) between male and female Dahl SS rats at 8 and 24 weeks on control and HF diet. Male and female rats at 8 weeks on diet showed no significant change in body weights between the control and HF diet fed groups (**Extended Data Figure 1)**. At 24 weeks, there was a modest increase in weight in both HF male and female animals compared to the control diet fed groups. Compared to their respective 8-week groups, MAP was found to be lower in the control diet fed rats but higher in the HF diet fed rats at 24 weeks in both males and females. This elevation in blood pressure in the HF diet fed Dahl SS rats was an expected result as previously documented^14^.

### Thoracic aortic PVAT is composed of distinct cell subtypes

Given that taPVAT is a complex tissue made up of varied cell types, we wanted to thoroughly examine cell type composition in our data. A total of twenty-five taPVAT samples from individual Dahl SS rats were used for library preparation and sequencing. Three individual samples were taken of each treatment group except for the 24-week HF diet fed female rat group which had four samples collected due to a sample clog resulting in a smaller number of recovered nuclei. In total, 83,849 nuclei were sequenced with a median of 3159 nuclei per sample. A median of 2501 transcripts and 1462 genes were detected per nucleus. Ambient RNA accounted for 4.1% of reads and was removed using SoupX^18^. Doublet detection and removal was performed with scDblFinder^19^ which labeled 6.1% of cells as doublets. After quality control and preprocessing, we had a remaining total of 71,813 cells and 20,743 genes for downstream analyses.

We subsequently performed Principal Component Analysis (PCA)^20^ and Uniform Manifold Approximation and Projection (UMAP)^21^ to reduce the dimensionality of the data for visualization. Cells were clustered using the Leiden algorithm^22^ at a resolution of 1.6 for subsequent cell-type annotation. The resulting cell clusters were annotated by examining the expression of marker genes for cell types previously reported in adipose tissue (**Figure 2A - 2B**)^23,24^. The initial “high-level” cluster annotation identified eight major cell types: adipocytes (*Plin1+*), endothelial cells (ECs, *Ptprb+*), fibroblasts (*Dcn*+), mesothelial cells (*Wt1*+), immune cells (*Ptprc*+), pericytes (*Rgs5*+), smooth muscle cells (SMCs, *Acta2*+), and neuronal cells (*Scn7a*+) (**Figure 2C**). Not surprisingly, taPVAT is primarily composed of adipocytes (64.1%), endothelial cells (16.0%) and fibroblasts (9.3%), with smaller populations of immune cells (4.7%), pericytes (3.4%), mesothelial cells (2.2%), smooth muscle cells (0.3%), and neuronal cells (0.2%) (**Figure 2D**). The substantial presence of endothelial cells highlights the richness of the PVAT microvasculature, as recently detailed by our group^25^.

**Figure 2:**
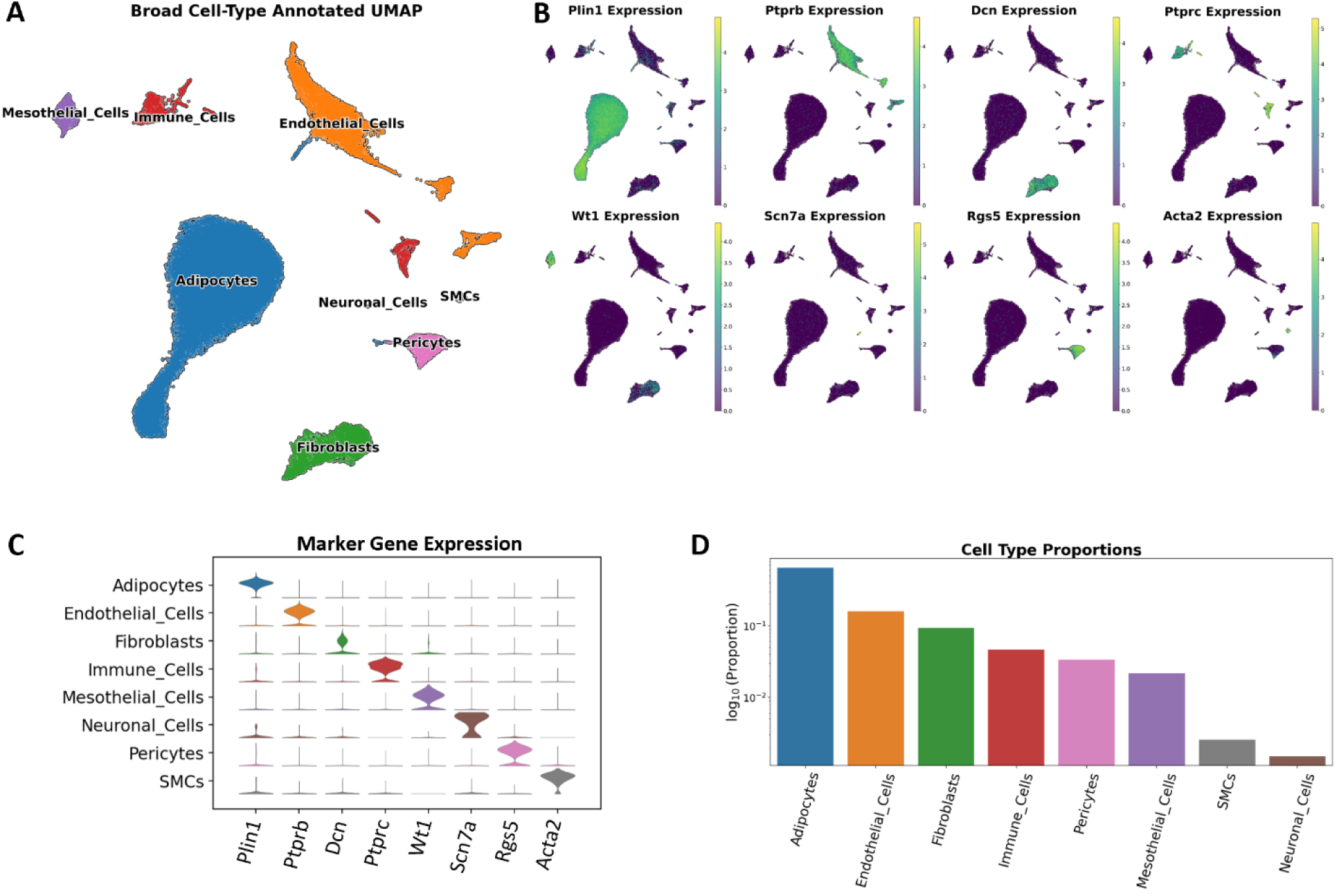
High level overview of cell types found in the dataset. **(A)** Broad, high-level, cell type annotated UMAP plot of all 71,813 cells inclusive of all samples. **(B)** Locations of marker gene expression on the UMAP. **(C)** Stacked violin plot of marker gene expression by high-level cell type annotation. **(D)** The proportion of each high-level cell type inclusive of all samples. Note that the y-axis is in log-scale.

We subsequently refined our cell type annotation by subclustering the immune cell, fibroblast, endothelial cell, and adipocyte populations (**Figure 3A - 3F**). The immune cell cluster separated into clearly defined groups of lymphocytes and myeloid cells. The lymphocytes were made up of T cells (*Skap1+,* 0.92%), B cells (*Pax5*+, 0.37%), and natural killer cells (*Gzmk+*, 0.34%). Likewise, myeloid cells (*Itgam+*) consisted of neutrophils (*S100a9+*, 0.22%), monocytes (*Fn1+,* 0.02%), M2-macrophages (*Cd163*+, 1.95%), M1-macrophages (*Lyz2*+, 0.45%), and dendritic cells (*Flt3+,* 0.14%) (**Figure 3B**). The fibroblast population is comprised of four subgroups based on the expression of *Bmper* and *Pi16*, similar to a previous study by Burl et.al^26^. The four subgroups were labeled Fibroblasts_Bmper+ (1.5%), Fibroblasts_Bmper+_Nrxn1+ (4.9%), Fibroblasts_Pi16+ (1.7%), and Fibroblasts_Pi16++ (1.1%) (**Figure 3C**). Similar populations of progenitor cells have been found in other adipose tissue cell atlases, however, there are variations in the naming convention of this cell type (i.e. adipocyte stem and progenitor cell (ASPC), fibro-adipogenic progenitors (FAPs), pre-adipocytes, etc.)^24,27–29^. Endothelial cells also had several subtypes (**Figure 3D**). We identified groups of endothelial cells expressing marker genes that have previously been associated with lymphatic (*Prox1*+, 1.5%), venous (*Vwf*+, 1.9%), arterial (*Fbln5*+, 3.2%) and capillary (*Kdr*+, 9.3%) endothelial cells^30,31^. Finally, we subset the adipocytes and found a large group of brown adipocytes marked by their expression of *Ucp1* that make up 57.4% of the overall cell population (**Figure 3E**). There were also two groups of *Pparg+Ucp1*- adipocytes that both express *Car3* but only one group expressed *Kcnip1*. The *Car3+Kcnip1+* cluster, annotated as Adipocytes_3 in the figures, and the *Car3+Kcnip-* cluster, annotated as Adipocytes_2, make up 4.2% and 2.5% of the cell population respectively. These groups may mark committed adipocyte progenitor cells (APCs), or they may be white or beige adipocytes^26,27^. The expression pattern of key marker genes showed clear segregation in these “low-level” cell type annotations (**Figure 3F**).

**Figure 3:**
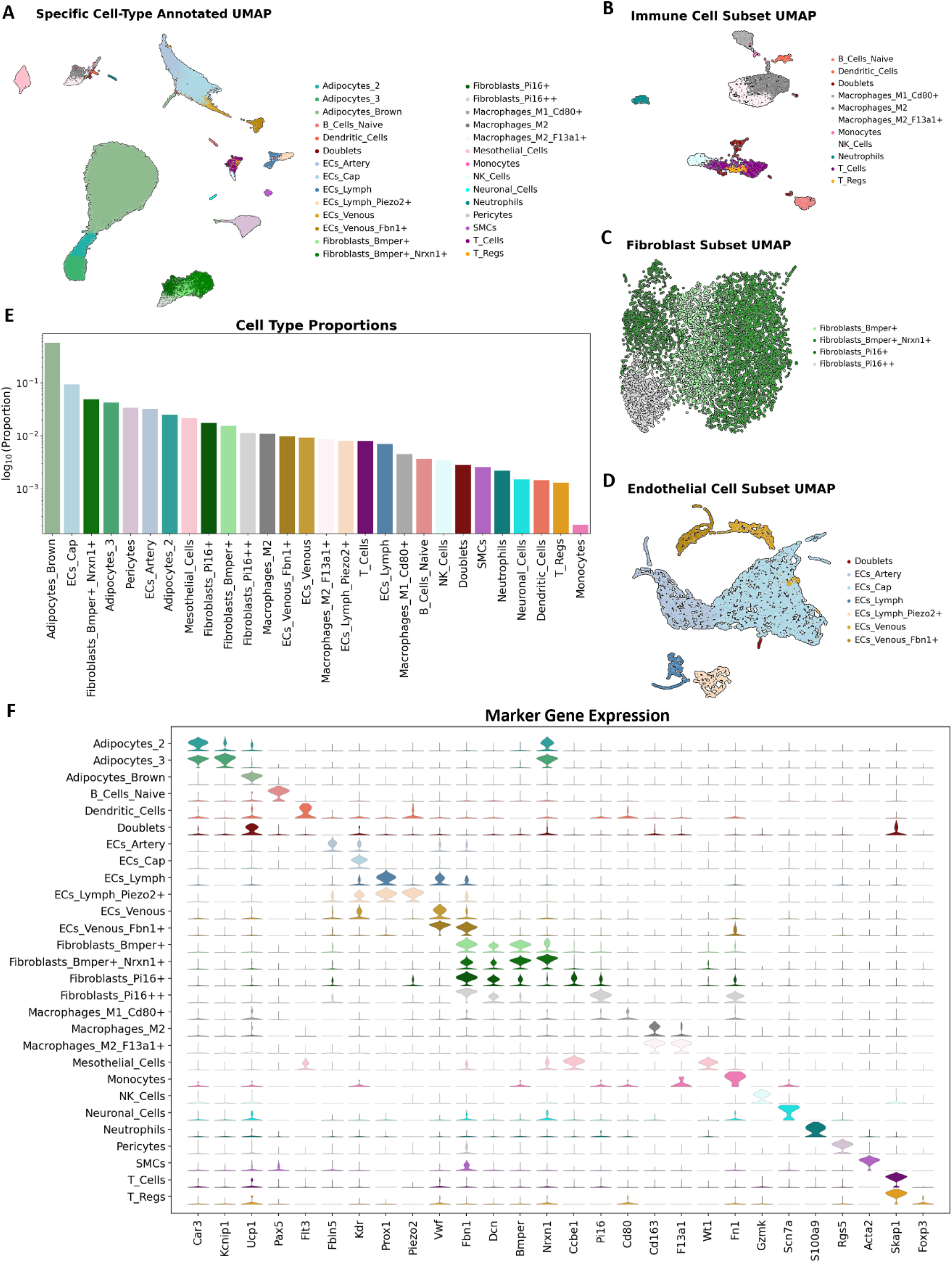
Low-level view of cell types found in the dataset. **(A)** UMAP of combined samples with specific, low-level, cell type annotations. **(B, C, D)** Subset and reprojection consisting of immune cells, endothelial cells, and fibroblasts. **(E)** The proportion of each low-level cell type inclusive of all samples. Note that the y-axis is in log scale. **(F)** Stacked violin plot of marker gene expression by low-level cell type annotation.

### Cell type proportions vary by diet, diet duration, and sex

The combined dataset with cell type annotations was split by sample into 25 subsets; and the respective cell type proportions were calculated for each individual sample. The samples were then grouped by treatment and cell type proportions were compared. A two-tailed Welch’s t-test was used to calculate significant differences in the cell type proportions across each treatment. The proportions varied between groups, especially when compared across time on different diets (**Figure 4A**). We initially considered the eight high-level, broadly identified cell types.

**Figure 4:**
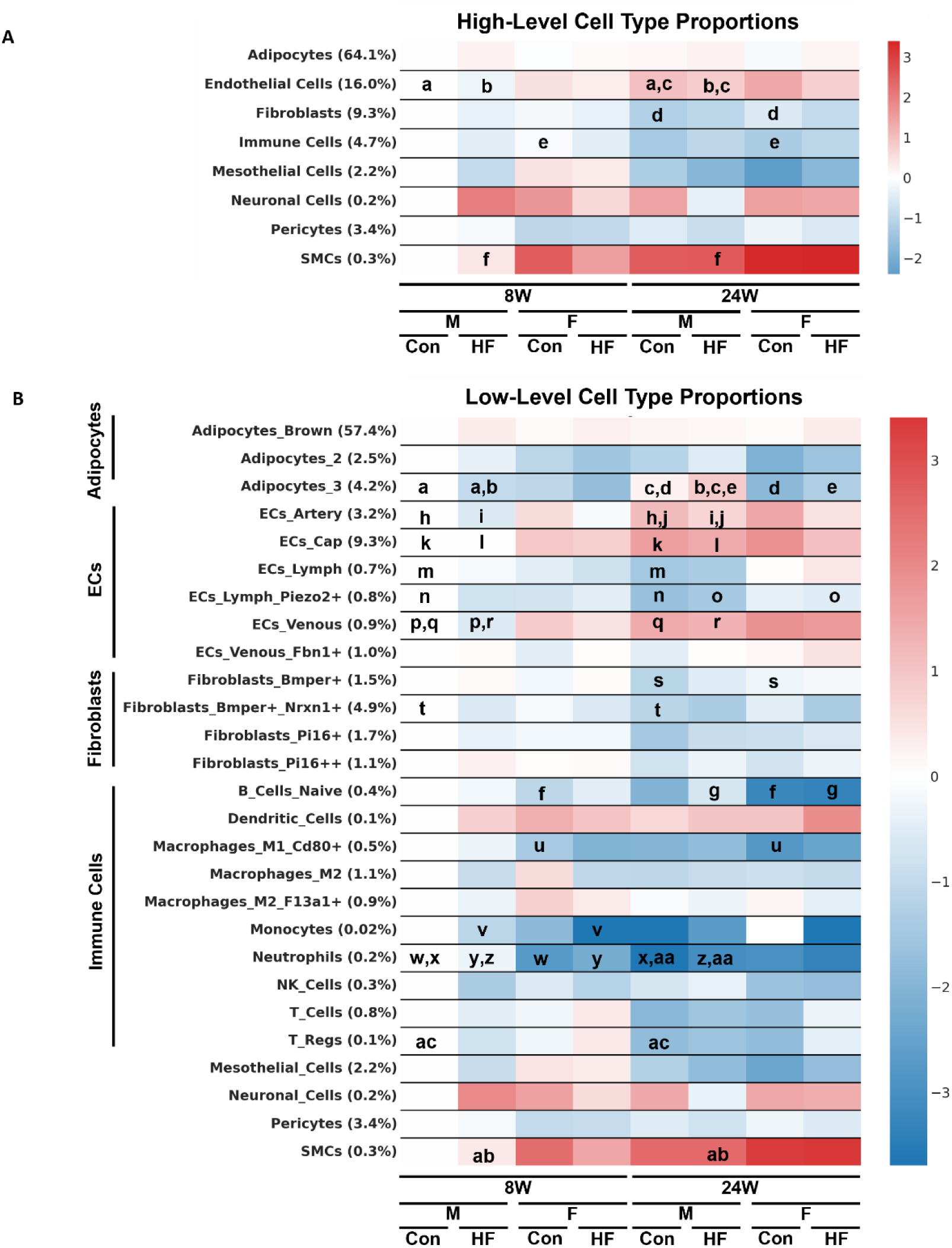
Heatmaps representing the proportions of (A) high-level and (B) low-level cell types and how they differ across each treatment category. The y-ticks are labeled with the cell types and their mean proportion across all treatment categories. The color bar legend indicates the log_2_(fold change) of the treatment proportion relative to the 8-week, control-diet fed, male rat treatment group for each cell type. All statistical comparisons were made using a two-tailed Welch’s t-test. Significant differences (p-value *≤* 0.05) between treatments are marked by the letters on the plot. I.e. “a” means that there is a significant difference in the proportion of endothelial cells between diet durations in male rats fed a control diet.

Interestingly, only endothelial cells showed differences in abundance between control and HF diet. Specifically, there was a significantly lower proportion of endothelial cells in the male rats fed a HF diet for 24 weeks compared to the control (**Fig. 4A [c]**). Between sexes, there was a significantly higher proportion of fibroblasts in the female versus male rats fed a control diet for 24 weeks (**Fig. 4A [d]**). This difference was not observed in the HF diet fed rats. Comparing the 8-and 24-week time points, we found a significantly higher proportion of endothelial cells at 24 weeks in male rats fed either a HF or control diet (**Fig. 4A [a,b]**), a lower population of immune cells at 24 weeks in female rats fed a control diet (**Fig. 4A [e]**), and a higher population of SMCs at 24 weeks in male rats fed a HF diet (**Fig. 4A [f]**).

We then tested cell type proportion differences in the low-level cell types and again found most differences occurred across diet durations (**Figure 4B**). First, we compared the HF diet-fed rats to the control diet-fed rats and found a smaller proportion of arterial ECs in male rats at 24 weeks (**Fig. 3B [j]**), a lower proportion of venous ECs in male rats at 8 weeks (**Fig. 4B [p]**), a smaller proportion of *Kcnip1+* adipocytes [Adipocytes_3] in male rats at 8-weeks (**Fig. 4B [a]**), a higher proportion of *Kcnip1+* adipocytes [Adipocytes_3] in male rats at 24 weeks (**Fig. 4B [c]**), and a higher proportion of neutrophils in male rats at 24 weeks (**Fig. 4B [aa]**).

Second, we analyzed the data for sex differences and found that the female rats had a lower proportions of *Kcnip1+* adipocytes [Adipocytes_3] at 24 weeks on both HF and control diets (**Fig. 4B [d,e]**), a higher proportion of *Bmper+* fibroblasts at 24 weeks on control diet (**Fig. 4B [s]**), a higher proportion of *Piezo2+* lymphatic ECs at 24 weeks on HF diet (**Fig. 4B [o]**), a lower proportion of neutrophils at 8 weeks in both HF and control diets (**Fig. 4B [w,y]**), a lower proportion of B cells at 24 weeks in the HF diet (**Fig. 4B [g]**), and a lower proportion of monocytes at 8 weeks on the HF diet (**Fig. 4B [v]**).

Finally, when comparing diet durations, we found that at 24 weeks there was: a lower proportion of *Bmper+Nrxn1+* fibroblasts in male rats fed a control diet (**Fig. 4B [t]**), a lower proportion of lymphatic ECs in male rats fed a control diet (**Fig. 4B [m]**), a lower proportion of *Piezo2+* lymphatic ECs in male rats fed a control diet (**Fig. 4B [n]**), and a greater proportion of arterial ECs in male rats fed either a HF or a control diet (**Fig. 4B [h,i]**). At 24 weeks there was also a higher proportion of capillary ECs in male rats fed a HF or control diet (**Fig. 4B [k,l]**), a higher proportion of venous ECs in male rats fed either a HF or control diet (**Fig. 4B [q,r]**), a greater proportion of *Kcnip1+* adipocytes [Adipocytes_3] in male rats fed a HF diet (**Fig. 4B [b]**), a higher proportion of SMCs in male rats fed a HF diet (**Fig. 4B [ab]**), a lower proportion of neutrophils in male rats fed either a HF or control diet (**Fig. 4B [x,z]**), a lower proportion of *Cd80+* M1-macrophages in female rats fed a control diet (**Fig. 4B [u]**), a lower proportion of B cells in female rats fed a control diet (**Fig. 4B [f]**), and a lower proportion of regulatory T cells in male rats fed a control diet (**Fig. 4B [ac]**).

### Cell type specific gene expression patterns are affected by diet, diet duration, and sex

We investigated the effect of three variables (diet, diet duration, and sex) across cell type on gene expression levels. We first studied gene expression differences across the whole tissue, inclusive of all cell types. When comparing all control diet samples to all HF diet samples (this includes 8-week, 24-week, male and female data), we found a total of 577 differentially expressed genes (DEGs) (**Figure 5A** top panel). The 304 upregulated DEGs in the HF diet were associated with carboxylic acid catabolic process and ion transport pathways (**Figure 5A** middle panel), whereas the 273 downregulated genes were linked to broad fatty acid metabolism (**Figure 5A** bottom panel). Similarly, when we compared the 8-week timepoint samples to the 24-week timepoint samples, we found 784 DEGs (**Figure 5B** top panel). The 258 upregulated genes at 24 weeks were largely associated with regulation of cell motility (**Figure 5B** middle panel), and the 526 downregulated genes were linked to immune responses (**Figure 5B** bottom panel). Lastly, comparing the male and female rat samples, we found 181 total DEGs (**Figure 5C** top panel). The 43 upregulated genes in female rats were associated with positive regulation of vesicle fusion pathways (**Figure 5C** middle panel), whereas the 138 downregulated genes were linked to leukocyte migration (**Figure 5C** bottom panel).

**Figure 5:**
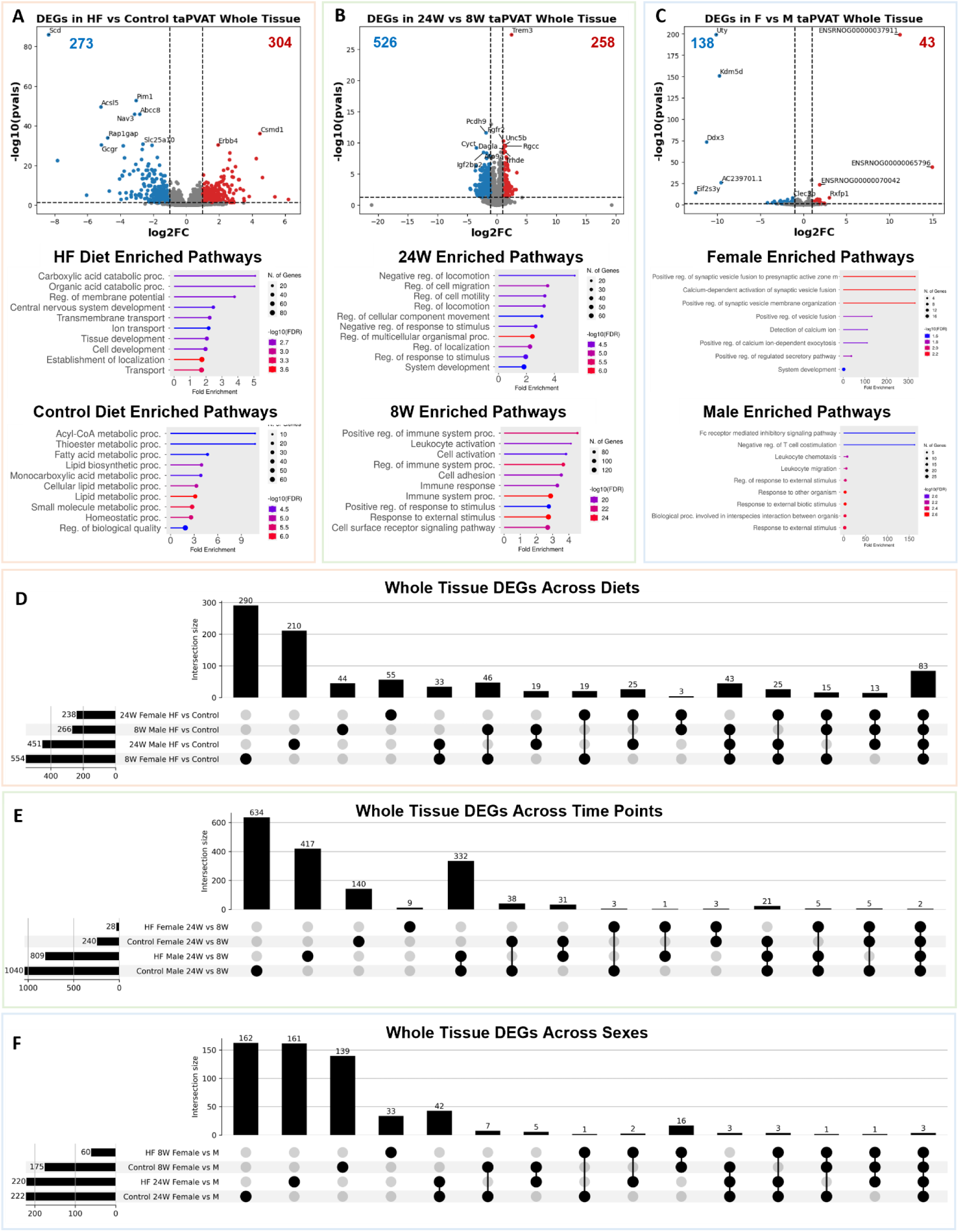
Examination of differential gene expression. **(A)** (Top) Volcano plot of differentially expressed genes (DEGs) between high fat and control diets. (Middle, Bottom) Enriched pathways in rats fed a HF or control diet (includes 8-week, 24-week, M and F data). **(B)** (Top) Volcano plot of differentially expressed genes (DEGs) between 8W and 24W diet durations. (Middle, Bottom) Enriched pathways in rats on diet for 24-weeks or 8-weeks (includes control diet, HF diet, M and F data). **(C)** (Top) Volcano plot of differentially expressed genes (DEGs) between female and male rats. (Middle, Bottom) Enriched pathways in female or male rats (includes control diet, HF diet, 8-week, and 24-week data). **(D)** Upset plot of DEGs across treatments in the whole tissue when comparing the control diet data to the HF diet data. There were 83 DEGs that were found in all 4 comparisons. **(E)** Upset plot of DEGs across treatments in the whole tissue when comparing the 8-week diet duration to the 24-week diet duration. **(F)** Upset plot of DEGs across treatments in the whole tissue when comparing the male and female rats.

When separating treatment groups by time on diet the number of DEGs in male rats fed a HF versus control diet rises from 266 at 8-weeks on diet to 451 at 24 weeks on diet. However, the number of DEGs in female rats fed a HF versus control diet falls from 554 at 8-weeks to 238 at 24-weeks. There are 83 DEGs conserved across the four treatment groups (**Figure 5D**). When comparing the 8- and 24-week time points, there are again more DEGs in the male rats than the female rats. The number of DEGs in both males and females is higher in control diet fed rats than in HF diet fed rats. There are only two DEGs conserved across the four treatment groups considering diet duration as a variable (**Figure 5E**). The fewest number of DEGs occurs when comparing female to male rats. The HF diet fed rats again had fewer DEGs than the control diet fed rats. The number of DEGs also increased in the 24-week on-diet group compared to the 8-week on-diet group. There were only three DEGs shared among all treatment groups (**Figure 5F**).

Building on the whole-tissue analyses, we conducted a cell-type specific differential gene expression analysis to dissect how each cell type individually responds to diet. This analysis highlighted cell-type-specific DEGs and enriched pathways that are otherwise masked in whole-tissue comparisons. When considering only the eight high-level cell types, adipocytes displayed the most pronounced gene expression changes across all three variables: diet, time on diet, and sex (**Extended Data Figure 2A-C**). The close overlap between adipocyte DEGs and whole-tissue DEGs implies that adipocytes are the primary driver of the observed whole-tissue gene expression changes. In adipocytes, HF diet upregulated DEGs were significantly enriched in tissue development and transport pathways. When comparing diets, downregulated genes in adipocytes were associated with broad fatty acid metabolism. After 24 weeks of diet exposure, adipocyte upregulated genes were linked to pathways involved in cell signaling and communication, while downregulated genes were associated with cell growth. Sex-specific analyses revealed enrichment for hormone synthesis and secretion pathways in males, whereas upregulated DEGs in females were associated with phosphorus metabolism.

Endothelial cells also demonstrated notable gene expression changes, particularly in response to the duration of diet (**Extended Data Figure 3A-C**). In 24-week versus 8-week diet comparisons, upregulated genes in endothelial cells were involved in fatty acid metabolism pathways and leukocyte activation. The downregulated genes were largely associated with cell-cell signaling pathways. This suggests a potential reduction in endothelial signaling with aging or prolonged dietary stress, coinciding with increased metabolic and inflammatory demands. Immune cells exhibited fewer DEGs across all three variables (**Extended Data Figure 4A-C**).

As with adipocytes and endothelial cells, genes downregulated in response to the HF diet in immune cells were significantly associated with broad fatty acid metabolism pathways. At 24 weeks on diet, upregulated genes were associated with fat cell differentiation, whereas the upregulated genes at 8 weeks were associated with somatic diversification and recombination of T-cell receptor genes. Female rats were highly enriched for fatty acid metabolism pathways, and male rats were enriched for immune cell response inhibition pathways. Pericytes, mesothelial cells, neuronal cells, and SMCs had low numbers of DEGs across all three variables. Lastly, when we consider the 28 low-level cell types, brown adipocytes had the highest number of DEGs across all comparisons (**Extended Data Figure 5A-C**). These findings collectively underscore the value of single-cell transcriptomics in uncovering cell-type-specific responses, which are otherwise masked in bulk tissue analyses.

### Potential early markers of hypertension in brown adipocytes

Adipocytes are typically omitted from single-cell studies due to their large size and buoyancy, which can interfere with droplet-based microfluidic barcoding systems. Using snRNAseq circumvents this issue. As a result, our dataset includes both stromal and adipocyte populations, with adipocytes comprising the majority of captured cells (64.1%). Brown adipocytes specifically represented just over half (57.4%) of all nuclei sequenced from taPVAT. There was not a significant difference in the proportion of brown adipocytes between diets, time on diets, or sexes. There were, however, many differences in the gene expression of this cell population.

Gene expression changes in brown adipocytes largely mirrored those observed when analyzing the whole tissue. This isn’t surprising given their dominant representation in the tissue. Of the 577 DEGs identified in the whole tissue between diets, 548 genes (95%) overlapped with DEGs specific to brown adipocytes (**Extended Data Figure 6A**). Given their abundance and the fact that they are the parenchymal cells of taPVAT, we focused subsequent gene expression analyses on the brown adipocyte population.

One of our main objectives was to identify changes in gene expression resulting from hypertension induced by a high fat diet. As previously stated, the Dahl SS rat is known to become hypertensive when fed a high fat diet, but only after being on diet for a sufficient period of time (∼16 weeks)^14^. Our time points of 8 and 24 weeks were specifically chosen so we could compare animals in a “prehypertensive” (8-week) vs. “hypertensive” (24-week) state (**Extended Data Figure 1**). To identify DEGs associated specifically with the hypertensive state, we compared HF versus control diet brown adipocytes at 24 weeks. We then made the same comparison at 8 weeks to uncover genes with consistent or diverging differential expressions. We flagged genes that were already dysregulated at 8 weeks and remained dysregulated at 24 weeks as potential early markers of hypertension. Genes were sorted by the magnitude of their fold changes and whether they were upregulated or downregulated at both 8 weeks and 24 weeks on diet. The top three early marker genes that were upregulated in the HF diet were *Csmd1*, *Sall3*, and *Obsl1*, and the top three downregulated genes were *Scd*, *Grk1*, and *Acsl5 (***Supplemental Data**). However, with only two timepoints, further validation of these potential early markers using intermediate timepoints will be required.

To further disentangle the effects of diet-induced hypertension (8 to 24 weeks on a HF diet) from normal aging (8 to 24 weeks on a control diet), we computed gene expression changes over time within each diet group. We first used DESeq2 to identify DEGs in brown adipocytes comparing across time on diet in rats fed the HF diet and rats fed the control diet. This returned two matrices of fold change values and adjusted *p-*values (Benjamini-Hochberg) for each gene comparing across 8-weeks and 24 weeks on diet, one for the control diet group and one for the HF diet group. Then, for each gene, we subtracted the absolute value of the control diet fold change from the absolute value of the high fat diet fold change, resulting in the set of differences in fold changes across diets from the 8-week to 24-week transition. We then identified the genes with maximal changes in the fold change. These genes change differently in rats that develop hypertension over time versus rats that experience normal aging.

Among the genes of interest, *Thsd4* emerged as a compelling candidate for further investigation. *Thsd4* expression increased as the rats aged in both diet groups. However, at 8 weeks on diet, brown adipocytes from HF-diet fed rats already showed strongly elevated *Thsd4* expression compared to the control diet fed group. Not only was the expression of *Thsd4* elevated in the HF group, but the proportion of brown adipocytes expressing the gene at all was much higher (∼13% in HF vs. ∼2% in control). This difference became even more pronounced at 24 weeks (∼32% in HF vs. ∼15% in control), suggesting early and sustained upregulation in response to high-fat diet exposure (**Extended Data Figure 6B**). *Thsd4* is involved in microfibril formation and encodes ADAMTSL6 which binds directly to fibrillin-1 and promotes fibrillin-1 matrix assembly^32^. *Thsd4* has not previously been strongly linked with blood pressure changes; however, decreased *Thsd4* expression has been associated with an increased risk for thoracic aortic aneurysm and dissection^32,33^. It is plausible that an increase in *Thsd4* expression causes an increase in fibrillin-1 matrix assembly which changes the composition of the PVAT extracellular matrix, potentially resulting in increased blood pressure. However, further studies are required to test this mechanistic hypothesis and validate *Thsd4* as an early biomarker of hypertension.

### Cell-cell communication analysis reveals alterations and heterogeneity in signaling between and within cell types

Given the multicellular composition of PVAT and its putative role in regulating vascular function in health and disease, we wanted to examine the complex web of cell-cell communication among the varied cell types in PVAT. For our preliminary analysis of communication among cell types in aggregate, we used NicheNet^34^, which uses single-cell gene expression and a prior model that incorporates intracellular signaling to characterize how ligands from a “sender” cell type functionally communicate with a “receiver” cell type, as reflected in induction of downstream target genes. We also used CellChat^35^ and CellPhoneDB^36^ within the LIANA^37^ framework to predict key ligand-receptor interactions driving condition-specific cellular responses.

There were broad similarities in inferred signaling pathways across sex and diet (**Table 1**). For example, vascular endothelial growth factor (VEGF) - Neuropilin-1 (NRP1) signaling (blue shading, **Table 1**) from adipocytes to endothelial cells was one of the top pathways in control male, control female, and HF female animals, but was absent in HF males. This is notable, given the known role of the VEGF - NRP1 pathway in regulation of angiogenesis, vascular remodeling, and permeability^38^. Conversely, SORBS1 – ITGA1 signaling (green shading, **Table 1**) from adipocytes to endothelial cells was one of the top interactions only in 24-week HF males. SORBS proteins are implicated in stiffness-sensing and contractile force generation^39^, suggesting an enhanced mechanical signal transduction role for adipocytes in PVAT under high-fat diet conditions. Likewise, multiple Thrombospondin-1 (THBS1) - integrin signaling pathways, which are involved in mechanotransduction, angiogenesis, and vascular remodeling^40,41^, were prominent in HF but not control 8-week males (**Extended Data Table 1**).

**Table 1:**
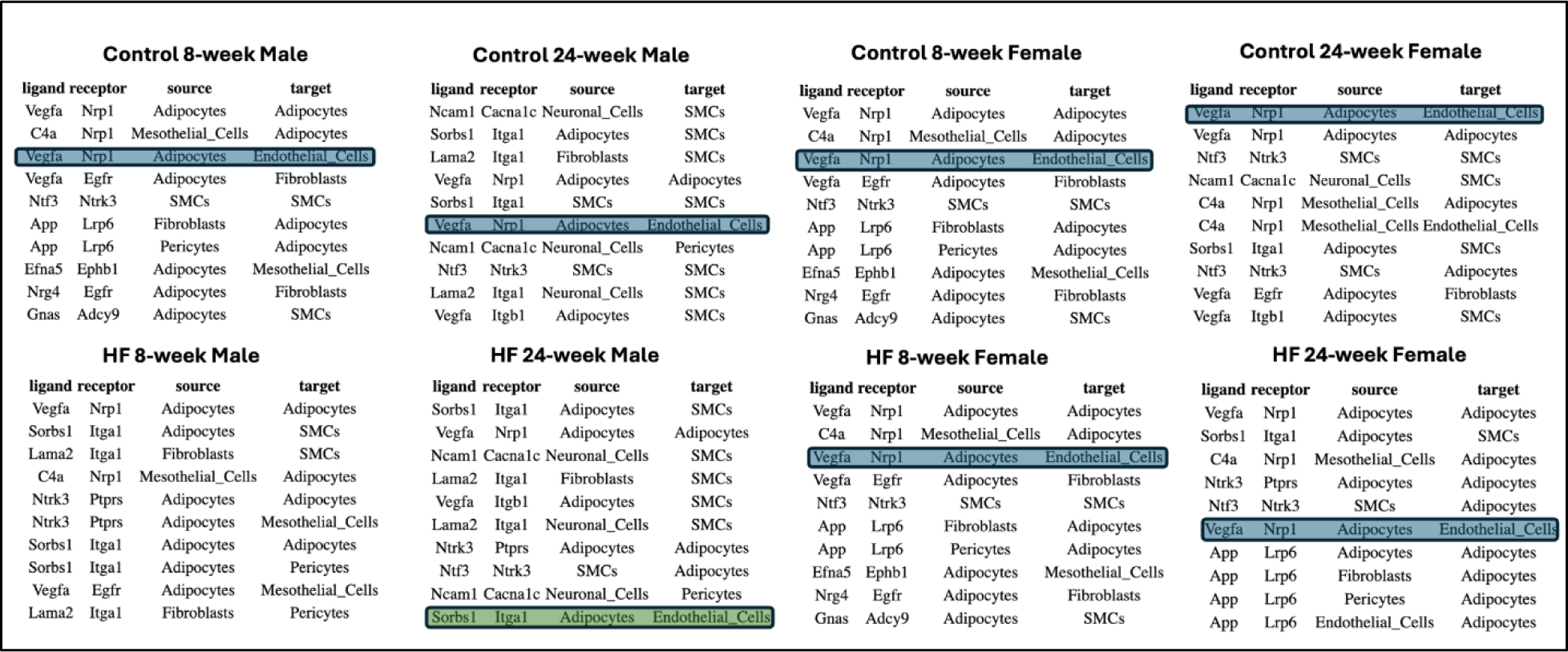
The most significant cell-cell communication pathways inferred across sex and diet. Vascular endothelial growth factor (VEGF) - Neuropilin-1 (NRP1) signaling (blue shading) from adipocytes to endothelial cells was one of the most active pathways in all groups except HF males. On the other hand, Sorbin and SH3 Domain Containing 1 (SORBS1) – Integrin Subunit Alpha 1 (ITGA1) signaling (green shading) from adipocytes to endothelial cells emerged as a notable intercellular signaling pathway only in 24-week HF males.

While the tools utilized above all infer interactions on a cell-population level, we wanted to delve deeper to investigate heterogeneity in signaling mechanisms within and between cell types at the level of individual cells. For this purpose, we utilized NICHES, a tool that infers cell-cell communication at a truly single-cell level based on simultaneous expression of ligands in the sending cell and cognate receptors in the receiving cell^42^. In control 8-week males, NICHES revealed distinct modules of signaling pathways involved in communication between specific sender and receiver cell types at single-cell level, with few interactions common to signaling among different cell type pairs (**Figure 6A**). For example, Adiponectin (ADIPOQ) to Adiponectin receptor ADIPOR2 signaling is largely restricted to adipocyte – adipocyte communication (**Figure 6A**, red highlight), while VEGF - vascular endothelial growth factor receptor 1 (FLT1) signaling was found in communication from both adipocytes and fibroblasts to endothelial cells (**Figure 6A**, green highlight). Using NICHES, we also projected individual cell-pair signaling events onto a 2-dimensional ligand-receptor interaction space, revealing distinct clusters of signaling events between cell-type pairs (**Figure 6B**). Zooming in to examine only adipocyte to endothelial cell communication, we found considerable heterogeneity in signaling mechanisms (pathways) between individual cell pairs, as reflected in distinct clusters of signaling events (**Figures 6C** and **6D**). Further, the occurrence of signaling pathways varied between control and HF diet (**Figure 6E**). For instance, TIMP3 – KDR signaling was less frequent and FGF1 – NRP1 signaling more frequent when comparing 24-week HF to 24-week control diet males (**Figure 6F**).

**Figure 6:**
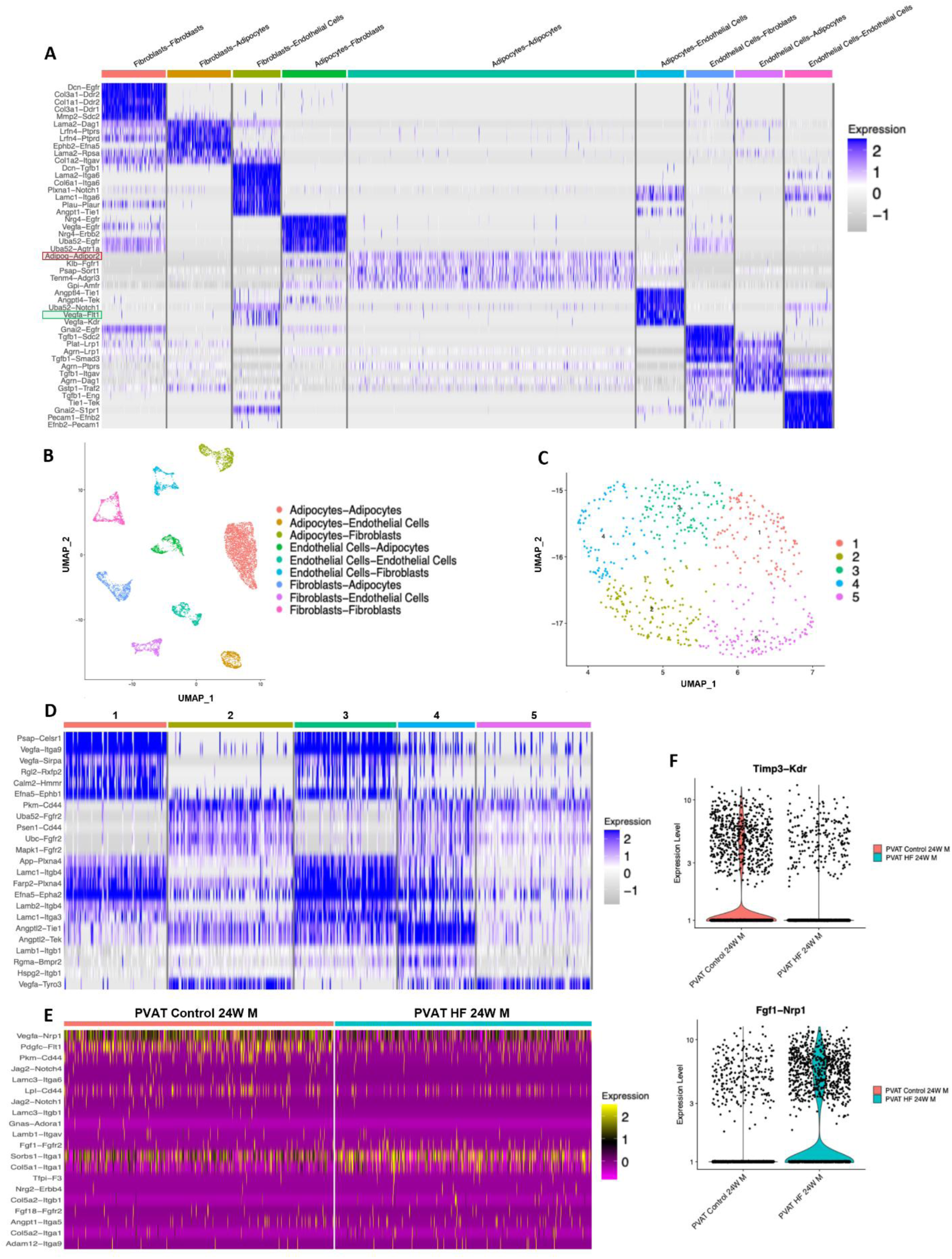
Cell-Cell Communication in PVAT. **(A)** Mapping of cell-cell interactions in PVAT from 8-week control males. **(B)** UMAP visualizations highlight distinct pairwise clusters of cell-cell interactions and communication patterns. **(C)** UMAP of adipocyte-to-endothelial cell interactions in 8-week control males. **(D)** Heatmap of ligand-receptor interactions between adipocytes and endothelial cells in 8-week control male, emphasizing key signaling pathways. **(E)** Comparison of top ligand-receptor interactions between adipocytes and endothelial cells in 24-week control and high-fat diet (HF) males. **(F)** Violin plots of *Timp3—Kdr* and *Fgf1—Nrp1*, showing increased activity in control conditions from 8 to 24 weeks and elevated expression in HF males over time.

### RNA velocity and pseudotime analysis reveal both gradual and abrupt transitions in gene expression dynamics depending on sex and time on HF diet

Next, we investigated gene expression dynamics in brown adipocytes, the predominant cell type in taPVAT. Specifically, we used scVelo^43^ to reveal transcriptional dynamics in response to a high-fat diet via generalized single-cell RNA velocity^44^. RNA velocity was calculated for each gene in each cell, following which the computed velocity vectors were embedded in a 2-dimensional diffusion map visualization for each group of brown adipocytes (**Figures 7A-D, top panels).** These visualizations show a clear directionality of state transition from control to HF diet in cells from both male and female animals. We also used scVelo to generate heatmaps of changes in adipocyte gene expression over latent time (pseudotime) from control to HF diet (**Figures 7A-D, bottom panels**). Only the top 500 most likely genes driving the pseudotime trajectory are shown, for animals at 8 weeks (**Figures 7A** and **7C**) and 24 weeks (**Figures 7B** and **7D**) on HF diet. The heatmaps show successive “waves” of expression of groups of driver genes along the transition trajectory. In most cases, the transition in adipocyte state appears to be gradual, as reflected in both the RNA velocity plots and heatmaps. However, in 24-week males (**Figure 7B**), we observe a more abrupt switch-like transition midway through the pseudotime trajectory, marked by a clear distinction between cells from control and HF-fed animals. The effects of diet on adipocyte state are thus sex-specific, especially in animals on the HF diet for a longer duration (24 hr.)

**Figure 7:**
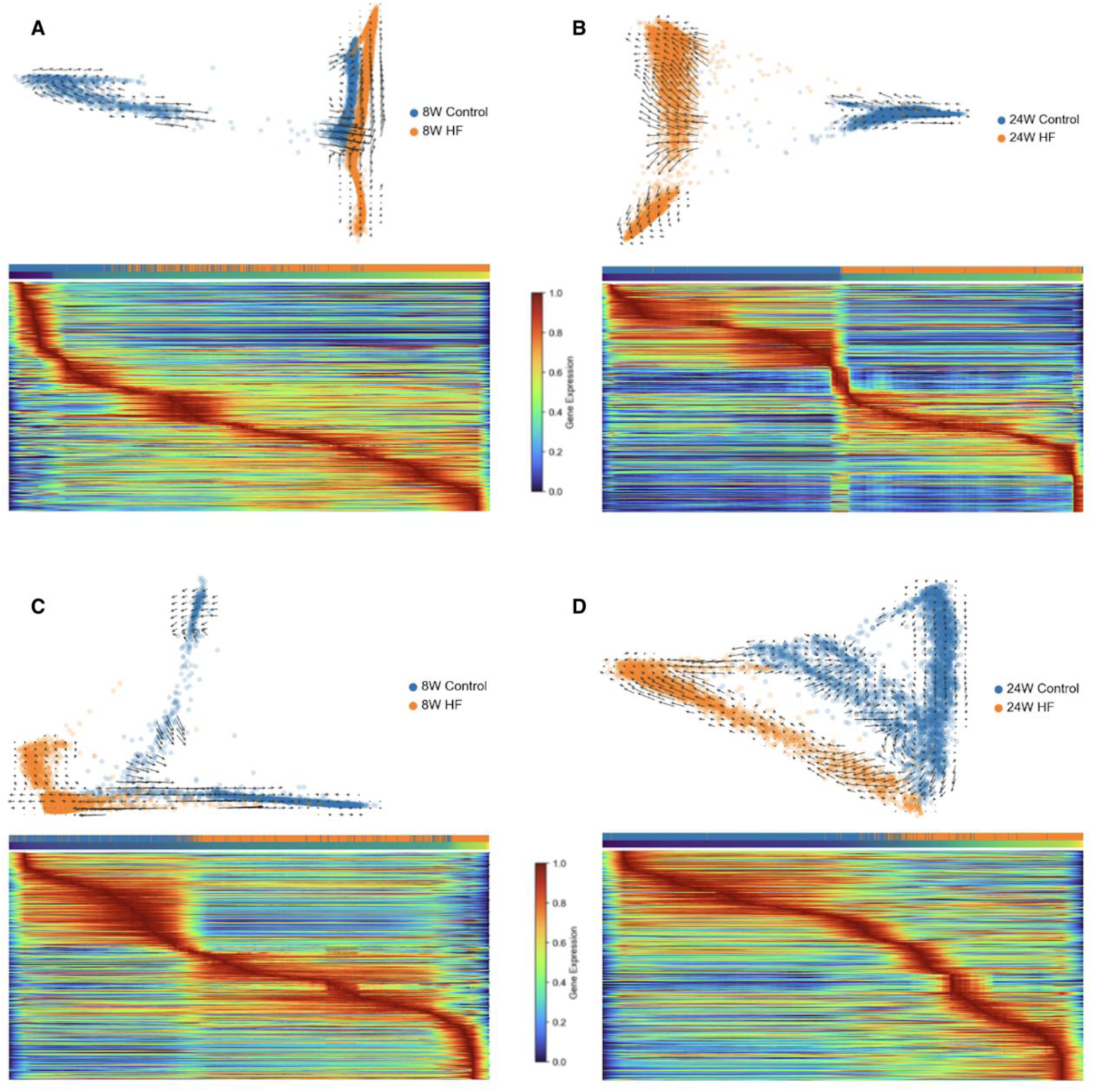
RNA velocity and latent-time gene expression heatmaps demonstrate progressively altered cell trajectories and gene programs in brown adipocytes. **(A)** 8-week control males versus 8-week high-fat males. The top panel shows RNA velocity vectors in a diffusion map embedding. The velocity vectors indicate the likely shift in gene expression states in nearby cells. Longer vectors indicate a greater RNA velocity (i.e., greater shift in gene expression space). The bottom panel depicts a heatmap displaying the gene expression magnitude of the top 500 genes driving the latent time trajectory. Red indicates high expression, while blue indicates low expression. Rows represent individual genes and columns correspond to cells, color coded by diet (blue for control and orange for high-fat). The progression through latent (pseudo) time is shown from purple (start) to yellow (end). **(B)** 24-week control males versus 24-week high-fat males. **(C)** 8-week control females versus 8-week high-fat females. **(D)** 24-week control females versus 24-week high-fat females.

### Gene regulatory network analysis identifies key transcriptional drivers of adipocyte state transition induced by HF diet

We then used CellOracle^16^ to infer key transcription factors (TF) that drive adipocytes towards a high-fat diet transcriptomic state by inferring condition-specific differential gene regulatory networks. Specifically, we wanted to identify TFs undergoing the largest functional change from control to high-fat diet conditions. TF functionality is defined per CellOracle by centrality or importance in a specific network using several alternative metrics: eigenvector centrality, out-degree centrality, and betweenness centrality. Higher eigenvector centrality measures characterize transcription factors that have more interactions with highly connected genes^45^. Out-degree centrality provides information about the number of genes TF regulates, while betweenness centrality gives an indication of which TFs act as a bottleneck between different highly connected sub-networks within a gene regulatory network^45–47^. Taking these three centrality measures together, we hypothesized that TFs that have a high score in each measure are likely to have greater functionality in regulating gene activity in a specific condition. The ratio of each centrality measure in HF vs. control diet thus characterizes TFs’ change in functionally between conditions. Higher ratios denote a TF that gained functionality in the high-fat diet group compared to the control-diet group. For each centrality ratio, TFs were ranked by magnitude of the ratio, TFs with the highest ratio receiving a rank of 0 and subsequent TFs receiving rank scores of 1, 2, and so on. Each TF’s “total score” was determined by summing the ranks from each measure. For example, in adipocytes from female animals at 24 weeks, the top ranked transcription factor was *Nr4a1* with a total ranked score of 3 due to its eigenvector, degree centrality out, and betweenness centrality ranks of 0, 1, and 2, respectively (**Extended Data Table 2**).

To validate the significance of these inferred transcription factors, we performed *in-silico* perturbation experiments using CellOracle. For each comparison, the top-ranked TFs, as determined by the combined centrality measurement scores (see **Methods**) were computationally knocked out from the gene regulatory network. This is depicted with a 2D UMAP embedding of adipocytes from: (i) 8-week females with knockout of *Npas2*, *Nr4a1*, and *Foxo3* (**Figure 8A**, upper panels); (ii) 24-week females with knockout of *Nr4a1*, *Epas1*, and *Nr4a2* (**Figure 8B**, upper panels); (iii) 8-week males with knockout of *Hlx*, *Bhlhe40*, and *Ezh2* (**Figure 8C**, top panels, and **Extended Data Figure 7)**; and (iv) 24-week males with knockout of *Cebpb*, *Nr4a3,* and *Nr4a2* (**Figure 8C**, bottom panels, and **Extended Data Figure 7.**

**Figure 8:**
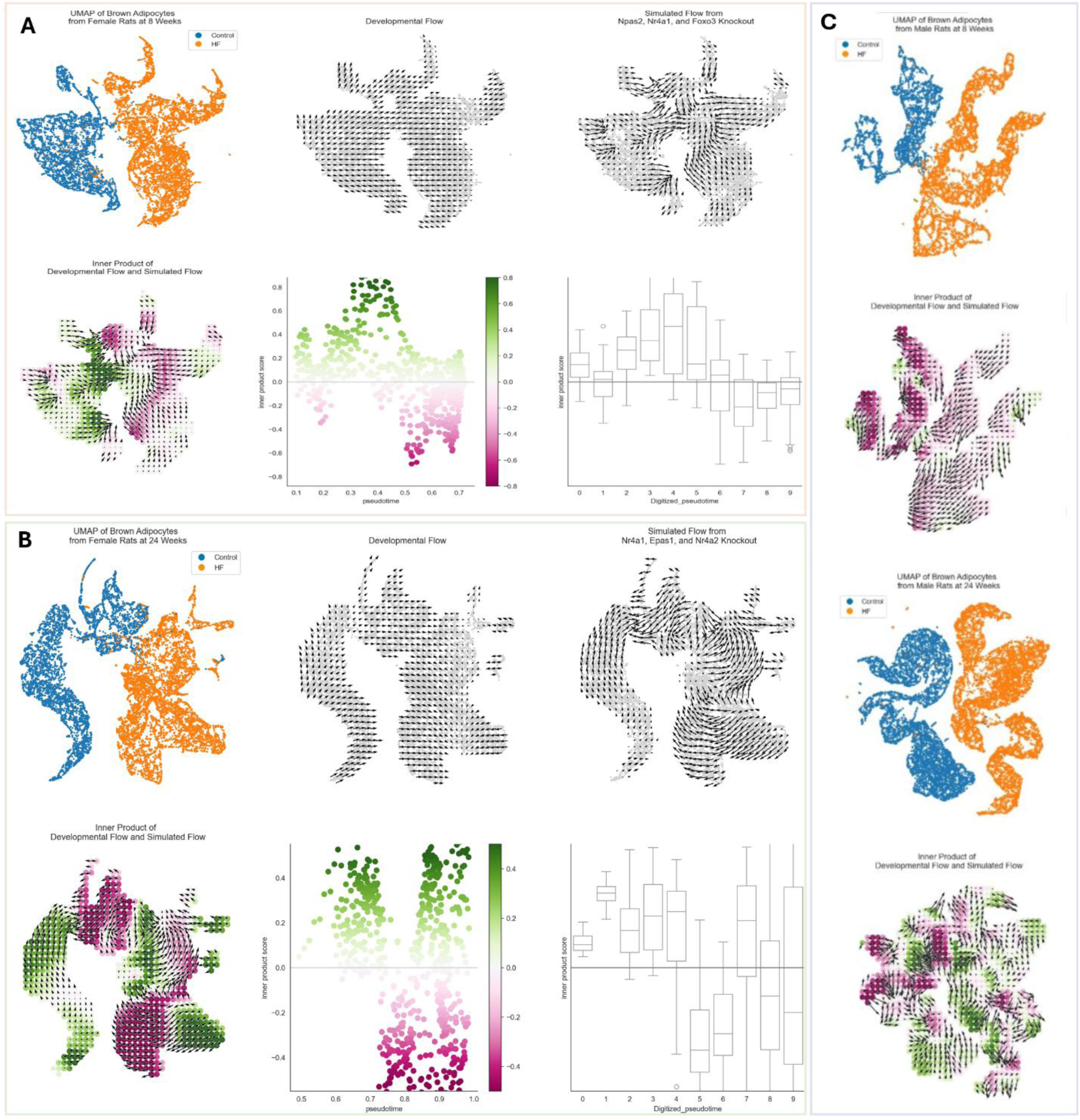
In–silico transcription factor knockout identifies potential regulators that either reinforce or resist diet-driven brown adipocyte state changes across cohorts. **A.** Panel A (top) depicts results from 8-week females (control versus high-fat diets). **B.** Panel B (bottom) depicts results from 24-week females (control versus high-fat diets). Top left: UMAP plot displaying control cells (blue) and high-fat diet cells (orange). Top center: Developmental flow vectors suggest a baseline progression from control cells to high-fat diet cells in gene expression space. Top right: Simulation vectors show the predicted transition in cell states after transcription factor knockout, representing the altered likelihood of a cell state change post-knockout. Bottom left: The dot product of developmental flow and simulation vectors, where green indicates a positive dot product, suggesting stabilization of the respective gene expression states, and pink indicates a negative dot product, implying resistance to transitions to these states. Bottom center: Scatter plot with each point representing a cell, colored by its dot product score. The x-axis represents pseudotime, and the y-axis quantifies the dot product score, illustrating gene expression state evolution over pseudotime. Bottom right: Box and whisker plots summarizing the data by binning cells into 10 groups across pseudotime, showing the distribution of dot product scores within each bin. **C.** Panel C depicts results between control and high-fat diets from 8-week males (top) and 24-week males (bottom). The top subfigures are UMAP plots displaying control cells (blue) and high-fat diet cells (orange). The bottom subfigures are the dot product of developmental flow and simulation vectors, where green indicates a positive dot product, suggesting stabilization of the respective gene expression states, and pink indicates a negative dot product, implying resistance to transitions to these states.

We show the results of our simulation for female rats (see **Extended Data Figure 7** for male animals). The “developmental / pseudotime vectors” (small arrows in **Figures 8A** and **8B**, upper middle panels) show the direction of diet-induced transition, while the “perturbation / knockout vectors” (small arrows in **Figures 8A** and **8B**, upper right panels) point in the direction of the most likely transition after simulated transcription factor knockout. The perturbation vectors demonstrate a uniform trajectory *away* from the high-fat diet-associated cell states, especially in the 8-week females. This is shown by a “perturbation score” (**Figures 8A** and **8B**, lower panels), the inner-product value between the pseudotime vectors and the knockout vectors. Green colors represent regions where the pseudotime vectors and perturbation vectors align in the same direction, such that these cell states become more stabilized. Conversely, pink colors represent cell states where differentiation and perturbation vectors are aligned in opposite directions, resulting in suppression of these states after knockout. Effectively, simulated knockout of top-ranked TFs disrupts the progression from control to high-fat diet associated cell states, even reversing it along parts of the pseudotime trajectory (**Figures 8A** and **8B**, lower middle and right panels), thus identifying driver TFs of cell state transition that could be potential therapeutic targets.

### Changes in gene expression induced by HF diet are coordinated and computationally predictable across cell types

Finally, we tested the hypothesis that cell-type-specific perturbations in gene expression can be modeled quantitatively as a function of their baseline gene expression state, allowing computational prediction of these perturbations. For this purpose, we used scVIDR^17^, a variational autoencoder-based generative deep learning model previously developed by us, to predict changes in gene expression levels in brown adipocytes derived from rats on the HF diet. For a given sex and timeframe (8- or 24-weeks), a model was trained on all cells except for the high-fat diet brown adipocytes. After training the model on either 8-week or 24-week gene expression, we predicted the expression of the top 100 differentially expressed genes and most highly variable genes in the brown adipocytes from either the 8-week or 24-week high-fat diet group. Specifically, a “diet perturbation-vector” in the latent space of the autoencoder was extracted based on the manifold hypothesis, which states that patterns in high-dimensional data, such as gene expression can be depicted with a lower-dimensional latent space. In this space, simple vector operations can capture complex behaviors in the original higher-dimensional space. We applied this idea to our data (see **METHODS** for details) by defining an HF-diet perturbation vector as the difference between the centroids of HF diet and control-diet cell clusters in latent space for each cell type. These perturbation vectors were then used to predict the high-fat diet gene expression for the brown adipocytes. Specifically, a linear regression model was fit to map each cell type’s control-diet centroid to its corresponding perturbation vector in order to predict the perturbation vector for the control-diet brown adipocytes. This perturbation vector was then added to the control brown adipocytes in latent space to predict the change in gene expression of brown adipocytes due to HF diet. The predictions were tested against the ground truth (actual data) gene expression for the top 100 differentially expressed genes (DEG) as well as all highly variable genes (HVG).

We obtained high prediction accuracy for all animal groups, with R² values ranging from 0.75 to 0.95 (**Figure 9**). For 24-week animals, prediction accuracy was consistently high: with R² = 0.89 (HVGs) and 0.94 (DEGs) for male animals and R² = 0.91 (HVGs) and 0.95 (DEGs) for females. 8-week males yielded R² values of 0.75 (HVGs) and 0.86 (DEGs), while females yielded R² = 0.87 (HVGs) and 0.86 (DEGs). These results suggest, remarkably, that there is an HF-diet-associated disease signature which characterizes perturbations in gene expression, and which can be predicted across cell type. This observation points to the viability of targeted drugs, or drug combinations, that could potentially reverse the altered gene expression profiles of multiple cell types involved in disease progression.

**Figure 9:**
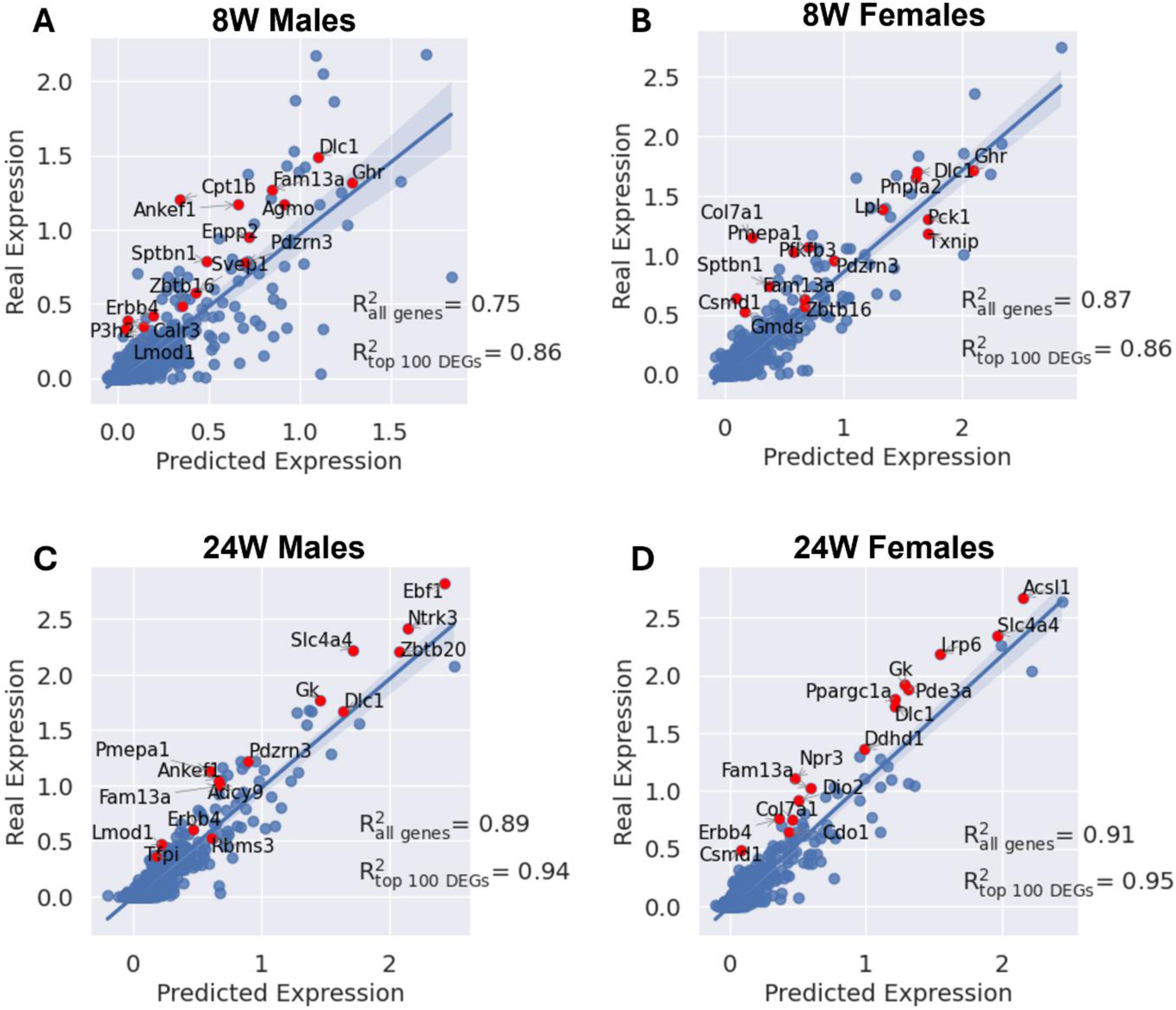
High-fat diet induced perturbation effects in brown adipocytes can be predicted from that of other cell types using latent-space computational modeling across cohorts. Prediction accuracy for high fat diet-associated brown adipocytes from **(A)** 8-week males, **(B)** 8-week females, **(C)** 24-week males, and **(D)** 24-week females. For each time and sex combination, the model was trained on the gene expression of all high-resolution cell types from both control and high-fat animal groups, except for the high-fat diet brown adipocytes. The altered gene expression in these cells was then predicted by adding the learned “disease signature” to the control brown adipocytes. Circles represent genes, with red circles marking the top fifteen differentially expressed genes in high-fat diet-associated brown adipocytes relative to control.

## DISCUSSION

This study presents a single-nucleus analysis of thoracic aortic perivascular adipose tissue (taPVAT) in Dahl SS rats, providing key insights into how diet, sex, and diet duration shape cell type composition, cell-type specific gene expression, and cell-cell communication. The data generated can serve as a rich resource for further study of this complex and fascinating tissue, and its role in vascular health and dysfunction.

We identified eight major cell types in taPVAT, predominantly adipocytes, endothelial cells, and fibroblasts, mirroring prior characterizations of adipose tissue^26,27^. However, the ability to further resolve these populations into twenty-eight lower-level subtypes, such as dendritic cells and lymphatic endothelial cells, offers novel insights. Notably, the predominance of brown adipocytes (57.4%) in this tissue reflects its specialized role in thermogenesis and metabolic regulation. Cell type composition changes highlighted significant effects of diet, diet duration, and sex. For instance, HF diet-fed male rats had a smaller proportion of endothelial cells, aligning with evidence of endothelial dysfunction under metabolic stress^48,49^. Differences between the sexes in fibroblast population include a higher proportion of *Bmper*+ fibroblasts in female animals. *Bmper* has recently been identified as a positive modulator of adipogenesis, thus the larger number of *Bmper*+ fibroblasts suggests that female rats may have a higher capacity to store excess lipid by creating new adipocytes^50^.

Differential gene expression analyses provided additional mechanistic insights. Adipocytes emerged as the primary driver of whole-tissue transcriptional changes, with upregulated genes enriched in pathways related to tissue development and transport under HF diet conditions. This observation suggests that adipocytes play a central role in adapting to metabolic challenges, likely through increased lipid storage and altered cell signaling. Endothelial cells also displayed significant gene expression changes, particularly with prolonged dietary exposure, highlighting their role in vascular inflammation and metabolic adaptation.

Our analysis of cell-cell communication pathways inferred from gene expression data revealed extensive sex-specific HF-diet induced alterations in ligand-receptor interactions between cell type pairs and signaling networks that may contribute to vascular dysfunction in hypertension. For example, VEGF-NRP1 signaling, linked to angiogenesis and vascular remodeling, was induced in high-fat females but repressed in high-fat males, suggesting sex-specific effects on vascular remodeling. In contrast, SORBS1-ITGA1 was increased in high-fat males, indicating enhanced mechanotransduction under dietary stress. Our analysis showed ADIPOQ-ADIPOR2 signaling to be limited to adipocyte autocrine communication, while VEGF-FLT1 facilitated adipocyte and fibroblast interactions with endothelial cells, highlighting diverse signaling modules in PVAT. Pathway-specific changes showed reduced TIMP3-KDR signaling, linked to extracellular matrix remodeling, and increased FGF1-NRP1 signaling under high-fat conditions, reflecting shifts in tissue remodeling processes. Clustering of ligand-receptor interactions at the level of individual cells revealed considerable heterogeneity of signaling even within the same cell-type pair, and distinct modules regulating localized vascular responses.

We should note that the varied computational tools used for cell-cell communication inference make somewhat different assumptions about ligand-receptor activity based on gene expression, which may not always reflect functional signaling activity. The discrepancies between tools like NicheNet, NICHES, and LIANA due to differing algorithms and databases highlight the need for cross-validation and experimental validation of predictions. The spatial organization of cells within PVAT, which can influence signaling patterns, was not directly assessed in this study. Validation of predicted signaling pathways using functional assays, such as ligand-receptor binding studies, would confirm our findings. However, the high-fat diet-induced changes in PVAT signaling we identified points to underlying mechanisms of vascular dysfunction and can suggest potential targets for mitigating PVAT-driven pathologies.

We also addressed the question: is cell state transition in response to high fat diet smooth and graded, or do gene expression profiles change more abruptly, suggestive of a switch-like transition at a critical threshold? Interestingly, while the diet-induced transition was gradual in general, male rats on HF diet for 24 weeks showed a more abrupt, switch-like transition as an effect of diet (**Figure 7**). Analyzing the adipocytes gene regulatory networks that could underlie these changes, we identified members of the nuclear receptor subfamily Nr4a (*Nr4a1*, *Nr4a2*, and *Nr4a3*) among the top three most functionally important transcription factors in every subgroup of rats except for 8-week males, where these receptors were found to be among the top 20 most important factors (**Extended Data Tables 2 and 3**). This suggests that the Nr4a subfamily is an important transcriptional regulator of high-fat diet associated hypertension and vascular dysfunction. Nra4 nuclear receptors are orphan receptors, and growing evidence implicates them in cardiovascular disease^51^. A reduction in adipogenic capacity is a major contributor to adipocyte hypertrophy which in turn drives several deleterious pathways and promotes inflammation^52^. Given that at least one member of the Nr4a subfamily (i.e., Nr4a1) is known to inhibit adipogenesis^53^, these transcription factors could be potential drug targets for addressing obesity-related hypertension. This possibility is supported by computational knockout experiments (**Figure 8** and **Extended Data Figure 7A** and **7B)** that demonstrated a suppression or potentially reversal, of high-fat diet associated cell states. One limitation of these analyses, however, is that we do not know which specific gene promoters are accessible for transcription factor binding due to lack of matched chromatin accessibility data, which would improve identification of target gene-specific upstream transcriptional regulators^16^

Finally, we used generative deep learning, specifically a variational autoencoder-based model that we have previously developed, to show that there is a consistent diet-associated perturbation signature that can be predicted across the multiple cell types making up PVAT. This is remarkable considering the diverse lineages of these cell types, and shows that the alteration in gene expression state of a given cell type in response to HF diet can be described quantitatively as a function of its baseline expression state. This finding can motivate the search for such coordinated transcriptomic perturbation signatures for other diseases and pathologies that manifest at the level of tissues made up of varied cell types. Such signatures could identify existing drugs with known transcriptomic perturbation profiles that could be repurposed to reverse specific disease signatures.

## METHODS

### Animal Models and Tissue Collection

Dahl SS male and female rats were purchased from Charles River Laboratory at 3 weeks of age. Animals were placed on control diet (10% kcal from fat; Research Diets, #D12450J) or high fat diet (60% kcal from fat; Research Diets, #D12492) upon arrival and housed in a 12:12 light dark light cycle for 8 or 24 weeks, with food and water available *ad libitum*. Use and care of animals complied with National Institutes of Health Guide for the Care and Use of Laboratory Animals (2011) and was approved by the MSU Institutional Animal Care and Use Committee (PROTO202000009). The study design and reporting follow the Animal Research: Reporting of In Vivo Experiments (ARRIVE) guidelines^54^. Due to the vital nature of adipose depots used in these snRNA-seq experiments, separate cohorts were utilized for the different time point tissue collections. Weights and MAPs for both the cohorts and specific rats used in the snRNA-seq experiments are therefore reported for transparency.

#### Blood Pressure measurements

Tail cuff plethysmography was used to measure blood pressure at the 8 and 24 week time point (CODA Non-Invasive blood pressure system, Kent Scientific, Torrington CA USA). Both systolic and diastolic pressures were measured, allowing for calculation of mean arterial pressure (MAP). At the same time, weights of the rats were recorded. We report BP and weights of the rats from which the taPVAT was taken for experimentation (below), along with the rest of their cohort, so that their context can be understood. Rats from each experimental group were selected for snRNA-seq using a computer-generated randomization scheme at the time of tissue collection to minimize allocation bias and ensure unbiased representation of each cohort.

At the conclusion of the study (8 or 24 weeks on diet), rats were anesthetized with isoflurane and a bilateral pneumothorax was created prior to dissection of the thoracic aorta. Under a stereomicroscope, a Silastic®-coated dish was placed on ice and filled with physiological salt solution [PSS, in mM: NaCl 130; KCl 4.7; KH_2_PO_4_ 1.18; MgSO_4_ • 7H_2_O 1.17; NaHCO_3_ 14.8; dextrose 5.5; CaNa_2_EDTA 0.03, CaCl_2_ 1.6 (pH 7.2)]. The taPVAT was removed from the vessel then snap frozen in liquid nitrogen and stored at –80° C.

### Nuclei Isolation and Flow Cytometry

Nuclei from frozen aortic perivascular adipose tissue were isolated with a modified protocol (dx.doi.org/10.17504/protocols.io.bkacksaw) of the Single Nuclei and Cytosol Isolation Kit for Adipose Tissues/Cultured Adipocytes (Invent Biotechnologies, #AN-029). Briefly, ∼150 mg of taPVAT and BAT were minced on a glass plate with small scissors and transferred to a sterile 1.5 mL microcentrifuge tube. On ice, samples were homogenized in N/C Buffer (from kit) using the supplied pestle. Samples were transferred to filter cartridges in collection tubes (from kit) and incubated, with the caps open, at −20°C for 30 minutes. Samples were centrifuged, filters discarded and tubes closed, vortexed, then centrifuged again. Supernatant was removed and nuclei were resuspended in Wash and Resuspension buffer (1X PBS with 1.0% BSA, 0.2 U/µL RNAse inhibitor) containing DAPI (10 µg/mL). Samples were filtered using a 40-um strainer and immediately sorted using a BD FACSAria IIu (BD Biosciences, San Jose, CA) at the MSU Pharmacology and Toxicology Flow Cytometry Core (facs.iq.msu.edu/), targeting 50,000 events/sample. Sorted nuclei were immediately processed with the 10X Chromium Next GEM Single Cell 3’ Reagent Kit v 3.1 (Dual Index) according to manufacturer’s recommended protocol (CG000315, Rev D), using the maximum sample volume of 43.2 µL. The generated cDNA was quantified, and size distribution determined using a D5000 ScreenTape assay on an Agilent 4200 TapeStation. Twenty-five percent of the total cDNA was used to generate the single nuclei 3’ gene expression libraries. Libraries were amplified with individual 10X Sample Indexes from the Dual Index Plate TT Set A according to manufacturer’s protocol, using a SimpliAmp Thermal Cycler. Cleaned up samples were eluted into 35.0 µL Buffer EB, with the average fragment size of the libraries determined using a D1000 ScreenTape assay on an Agilent 4200 TapeStation.

### Library Quantification

The KAPA Library Quantification Kit was used to quantify library samples, with samples diluted 1:1000, 1:5000, 1:10000, and 1:20000 in DNA Dilution Buffer (10 mM Tris-HCl, pH 8.0 −8.5 + 0.05% Tween 20). Six µL KAPA SYBR FAST qPCR Master mix and 4 µL of 1) no template control (RNase-free water), 2) each diluted sample, or 3) kit supplied DNA Standard were pipetted into a standard 96-well PCR plate, in triplicate, and the plate covered with an adhesive film to prevent sample evaporation. The plate was centrifuged for 1 minute at 1000 rpm in an Eppendorf 5804 centrifuge and placed in a QuantStudio 6 Flex Real-Time PCR System. The following conditions were run: 1 cycle of 95°C 5 minutes; 35 cycles of 95°C 30 seconds, 60°C 45 seconds, followed by a standard melt curve (to ensure the lack of primer dimers). The KAPA Library Quantification Data Analysis Template was utilized to determine the concentration of the undiluted libraries, which were submitted to Novogene for 150bp paired-end sequencing at a target depth of 50,000 reads/nuclei on a NovaSeq6000.

### Computational Analysis of Single-cell Data

The quality of the raw reads was initially checked using FastQC v0.11.7. Then, sequencing files were aligned to the rat reference genome (Rnor 7.2) using the 10X Genomics Cell Ranger v.7.1.0 pipeline^55^. The resulting h5ad files of aligned reads were then preprocessed with the R packages SoupX^18^ and scDblFinder^19^ to remove ambient RNA and doublets respectively. The parameters used for initial quality control before running SoupX and scDblFinder were the same as outlined in the “Single Cell Best Practices” manual^56^ except for the cut off for the percentage of mitochondrial gene counts. We used a 3% cut-off instead of a 10% cut-off since we had lower mitochondrial counts due to our use of single-nuclei sequencing instead of single-cell. SnRNA-seq is expected to exclude the sources for mitochondrial RNA. Individual samples were further preprocessed using the scanpy package^57^ with cutoffs for the minimum number of genes per cell and minimum number of cells per gene. Genes found in 3 or fewer cells were removed, as were cells with fewer than 100 measured transcripts. Cells that had greater than 15000 reads or 4500 measured genes were also discarded. The package scDblFinder gives each cell a score between 0 and 1 based on how likely it thinks that cell is a doublet. Cells assigned a 1 are automatically labeled as doublets, to be more conservative about our approach, cells with a score of 0.5 or greater were removed. After these quality control steps, the anndata objects for each sample were combined, and additional preprocessing with scanpy was done. Cells that were outside of 5 median absolute deviations from the medians of the log1p_n_genes_by_counts and log1p_total_counts were labeled outliers and discarded. The high-dimensional data was then visualized using principal component analysis (PCA)^20^ and Uniform Approximation and Projection (UMAP)^21^. Batch integration was performed with scvi-tools^58^ to remove technical noise from the reduced-dimension plots. Then clusters were assigned using the Leiden algorithm. We had an informed understanding of the types of cells that we might find thanks to our previous work on building a cell atlas of rat taPVAT^23^. We labelled Leiden clusters with cell types based on the expression of marker genes that had been previously reported to be found in adipose tissue (**Figure 2A - 2B**). Cell type annotation was first done at a high-level (i.e. immune cells) to identify major cell types, then refined via subclustering to identify lower level cell types (i.e. macrophages). Differential gene expression (DEG) analysis was done with biological replicates using the DESeq2 package implemented in python^59,60^. Upset and volcano plots were generated using a custom Python script available in this project’s GitHub repository. ShinyGO v0.81^61^ was used to perform pathway enrichment analysis, specifically, overrepresentation analysis. This calculated pathways in the Gene Ontology biological process 2023 database^62^ that the DEGs were associated with. Significantly upregulated DEGs (*log2fold change≥1 and p≤0.05)* were input to ShinyGO separately from significantly downregulated DEGs (*log2fold change≤−1 and p≤0.05)*.

### Cell-cell Communication analysis

We used NicheNet to analyze cell-cell communication among the different cell types within PVAT, predicting how ligands from sender cells influence target gene expression in receiver cells. NicheNet integrates expression data with a model of signaling and gene regulatory networks, constructing a weighted network of ligand-receptor, intracellular signaling, and gene regulatory interactions. This approach employs a Personalized PageRank algorithm to compute ligand-target regulatory potential scores, prioritizing ligands based on their ability to predict differentially expressed genes in receiver cells. Potential signaling pathways are inferred with model parameters optimized using Bayesian methods. Our analysis focused on the top 700 highly expressed genes to ensure robust identification of ligand-receptor-target gene interactions.

We used NICHES to map cell-cell interactions at a single cell level focusing on ligand receptor signaling within different groups. NICHES constructs an adjacency matrix to define potential cell connections and builds matrices to capture ligand receptor signaling, the effect of individual cells on each other, the overall system’s influence on a cell, and each cell’s impact on its surroundings. We then perform differential analysis to identify unique pathways within cellular clusters and uncover fine grained signaling patterns.

We also applied the LIgand-receptor ANAlysis Framework (LIANA) to examine communication patterns. LIANA has functionalities to use tools like CellChat and CellPhoneDB to infer cell-cell communication by analyzing paired ligand-receptor expression across different cell types. CellChat builds on a ligand-receptor interaction database (CellChatDB) derived from KEGG^63,64^ and recent literature, identifying differentially expressed signaling genes and calculating intercellular communication probabilities using a mass action model and random walk propagation. CellPhoneDB uses compiled database of ligand-receptor interactions and integrates it with the expression data to find meaningful interactions between cells. Both the tools use p-values to determine statistically significant interactions.

### RNA velocity analysis

To obtain the RNA velocity for each gene in each cell and to create the velocity vector embedding figures, we used the standard scVelo workflow, with minor changes. We used velocyto^44^ to generate the unspliced and spliced RNA count matrices that scVelo requires to calculate RNA velocity. We provided velocyto with (1) a filtered list of barcodes that were obtained from CellRanger’s cell calling algorithm, (2) bam files for each rat that were also generated with CellRanger, and (3) the rat reference genome annotation file (gtf) obtained from Ensembl (Rnor 7.2).

Velocyto outputs a loom file that includes unspliced and spliced reads for each rat. The biological replicates from each of the eight experimental conditions were then merged using loompy^65^, a standard package recommended by velocyto’s workflow, resulting in eight loom files. Depending on the analysis, sex- and diet-specific loom files were merged with the main single nuclei RNA sequencing dataset, using scVelo’s built in functions. The dataset, now containing unspliced and spliced reads, was filtered to include only brown adipocytes with at least 100 genes and for genes that were present in at least three cells. The dataset was then processed and analyzed with scVelo’s dynamical modeling workflow to generate the RNA velocity plots and heatmaps.

Diffusion maps were constructed using Scanpy methods (i.e., sc.tl.diffmap) and RNA velocity vectors were plotted on the diffusion map embeddings.

### Gene Regulatory Network Analysis

To obtain gene regulatory networks (GRN) and perform simulated knockouts, we used the standard CellOracle^16^ workflow. The dataset was preprocessed according to CellOracle’s standard workflow and brown adipocytes were selected. Since no scATACseq data was available, we used CellOracle’s built-in rat promoter base gene regulatory network, which provides baseline information for gene regulatory networks (GRNs) in rats. Using this built-in base promoter GRN, CellOracle builds a GRN model specific to our dataset based on user-defined clusters and calculates various network scores for transcription factors. For each analysis, GRNs were constructed for each condition (“time diet” cluster). For example, for each sex, 8-week control diet and 8-week high-fat diet groups would each have a unique GRN.

To identify transcription factors of possible importance, we used information from three graph centrality measures: (1) betweenness centrality, (2) eigenvector centrality, and (3) degree-out centrality. In each analysis (females or males, 8- or 24-weeks) the ratio of graph measures in the high-fat diet versus control diet was calculated for each transcription factor. Larger ratios mean that the specific graph centrality measure became more prominent in the high-fat diet group compared to the control diet group for that specific transcription factor. Graph centrality ratios are then sorted from largest to smallest and ranked by position, starting from zero. Larger ratios for each centrality measure, therefore, correspond to smaller rank values. Finally, the rank values for the three centrality measures are summed for each transcription factor. This summation is sorted from smallest to largest, and the three transcription factors with the smallest total value are chosen to be computationally knocked out. The combination of betweenness centrality, eigenvector centrality, and degree-out centrality suggests these transcription factors may be highly influential within the gene regulatory network and in enabling the transition from control diet associated gene expression states to HF diet gene expression states.

### Prediction of gene expression perturbation using scVIDR

For full details on the scVIDR method, we refer the reader to Kana et al., 2023^17^. In brief, scVIDR is a variational autoencoder model which first encodes high dimensional data (thousands of genes) into a lower dimensional latent space (100 dimensions). The latent space allows for simplified mathematical operations on the latent representations of the cells. The latent representations are then fed through a decoder which reconstructs the full dimensionality of the original data. Before variational autoencoders can be useful, the encoder and decoder must be trained. The model was trained in a leave-one-out paradigm, where all cell types except for the cell type to be predicted (i.e., brown adipocytes from the HF diet rat being assessed) were used during training. A perturbation vector for the HF diet condition was computed in the latent space as the difference between the mean latent representation of control and HF diet cells, calculated separately for each cell type. A linear regression model was then fit to map each cell type’s control latent representation to its corresponding perturbation vector.

To predict the HF diet response in brown adipocytes, the model estimated a perturbation vector using the mean latent representation of control-fed brown adipocytes. This predicted perturbation vector was added to the control latent representation to generate a latent prediction of the HF diet condition, which was then decoded to produce a gene expression matrix. To assess the model’s predictive performance, the predicted gene expression matrix was compared to the ground-truth 24-week HF diet brown adipocyte data to assess model performance.

## Data and code availability

Processed data can be conveniently queried through our implementation of the CellxGene Annotate web application located here: https://pvatcellatlas.azurewebsites.net/. Raw sequencing files are privately available for reviewers upon request and will be made publicly available through GEO upon publication. The GitHub repository for this project containing all notebooks and code is also available for reviewers upon request and will be made publicly available upon publication.

## Acknowledgements

This work was supported by the National Heart, Lung, and Blood Institute (NHLBI) P01HL152951, the National Institute of Environmental Health Sciences of the National Institutes of Health (NIEHS) R01 ES031937, and the AgBioResearch at Michigan State University with funding from the Hatch Act capacity funding program (Accession Number 1018334) from the USDA National Institute of Food and Agriculture. This work was also supported in part through computational resources and services provided by the Institute for Cyber-Enabled Research at Michigan State University. The content is solely the responsibility of the authors and does not necessarily represent the official views of the National Institutes of Health.

## Extended Data

**Extended Data Table 1:**
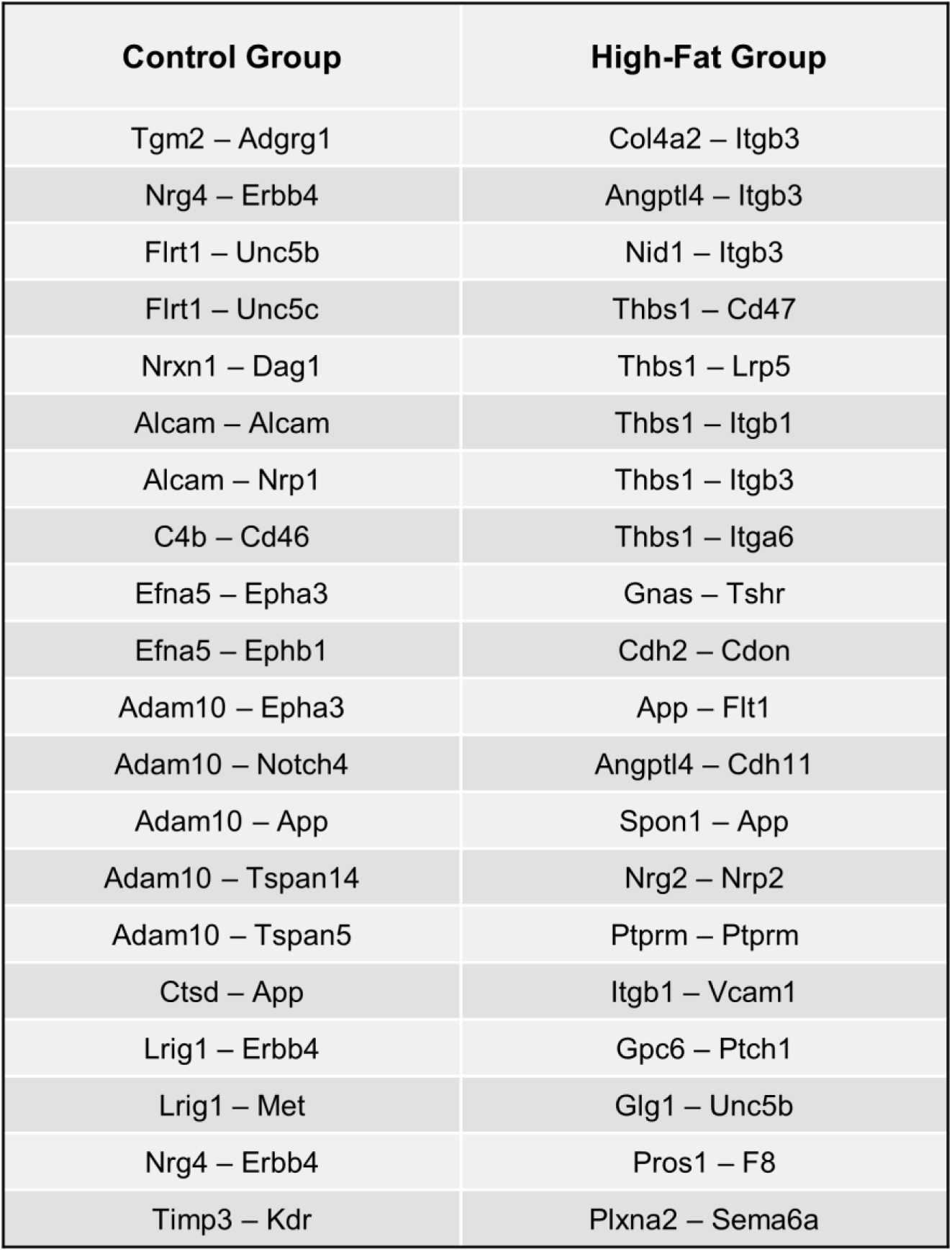
Top common interactions between adipocytes and endothelial cells in 8-week male rats, specific to control and high-fat diet groups, identified across all cell-cell communication tools.

| Control Group | High-Fat Group |
| --- | --- |
| Tgm2 – Adgrg1 | Col4a2 – Itgb3 |
| Nrg4 – Erbb4 | Angptl4 – Itgb3 |
| Flrt1 – Unc5b | Nid1 – Itgb3 |
| Flrt1 – Unc5c | Thbs1 – Cd47 |
| Nrxn1 – Dag1 | Thbs1 – Lrp5 |
| Alcam – Alcam | Thbs1 – Itgb1 |
| Alcam – Nrp1 | Thbs1 – Itgb3 |
| C4b – Cd46 | Thbs1 – Itga6 |
| Efna5 – Epha3 | Gnas – Tshr |
| Efna5 – Ephb1 | Cdh2 – Cdon |
| Adam10 – Epha3 | App – Flt1 |
| Adam10 – Notch4 | Angptl4 – Cdh11 |
| Adam10 – App | Spon1 – App |
| Adam10 – Tspan14 | Nrg2 – Nrp2 |
| Adam10 – Tspan5 | Ptprm – Ptprm |
| Ctsd – App | Itgb1 – Vcam1 |
| Lrig1 – Erbb4 | Gpc6 – Ptch1 |
| Lrig1 – Met | Glg1 – Unc5b |
| Nrg4 – Erbb4 | Pros1 – F8 |
| Timp3 – Kdr | Plxna2 – Sema6a |

**Extended Data Table 2:**
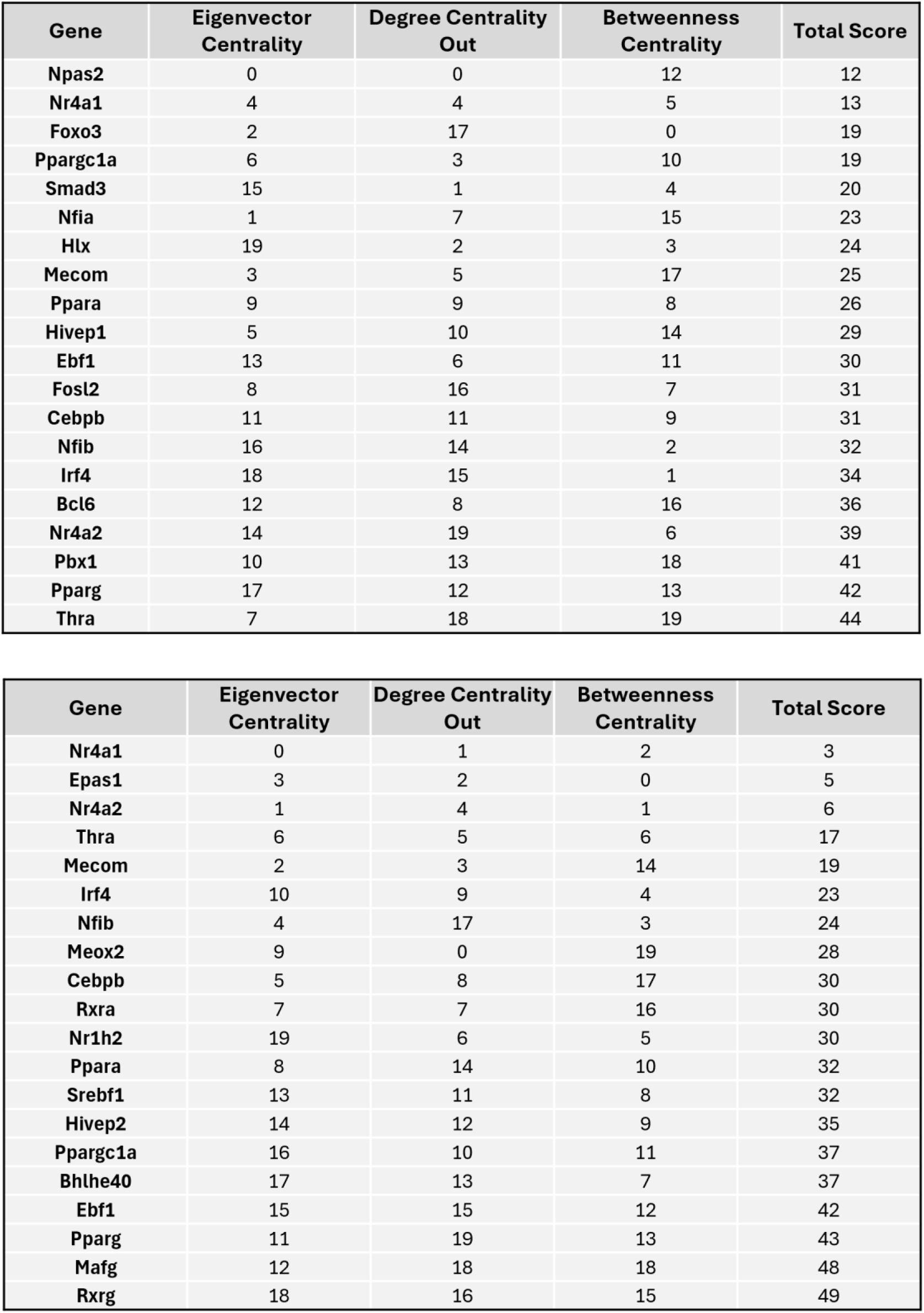
Top transcription factors identified by CellOracle for Females (8W top, 24W bottom)

| Gene | Eigenvector Centrality | Degree Centrality Out | Betweenness Centrality | Total Score |
| --- | --- | --- | --- | --- |
| Npas2 | 0 | 0 | 12 | 12 |
| Nr4a1 | 4 | 4 | 5 | 13 |
| Foxo3 | 2 | 17 | 0 | 19 |
| Ppargc1a | 6 | 3 | 10 | 19 |
| Smad3 | 15 | 1 | 4 | 20 |
| Nfia | 1 | 7 | 15 | 23 |
| Hlx | 19 | 2 | 3 | 24 |
| Mecom | 3 | 5 | 17 | 25 |
| Ppara | 9 | 9 | 8 | 26 |
| Hivep1 | 5 | 10 | 14 | 29 |
| Ebf1 | 13 | 6 | 11 | 30 |
| Fosl2 | 8 | 16 | 7 | 31 |
| Cebpb | 11 | 11 | 9 | 31 |
| Nfib | 16 | 14 | 2 | 32 |
| Irf4 | 18 | 15 | 1 | 34 |
| Bcl6 | 12 | 8 | 16 | 36 |
| Nr4a2 | 14 | 19 | 6 | 39 |
| Pbx1 | 10 | 13 | 18 | 41 |
| Pparg | 17 | 12 | 13 | 42 |
| Thra | 7 | 18 | 19 | 44 |
| Nr4a1 | 0 | 1 | 2 | 3 |
| Epas1 | 3 | 2 | 0 | 5 |
| Nr4a2 | 1 | 4 | 1 | 6 |
| Thra | 6 | 5 | 6 | 17 |
| Mecom | 2 | 3 | 14 | 19 |
| Irf4 | 10 | 9 | 4 | 23 |
| Nfib | 4 | 17 | 3 | 24 |
| Meox2 | 9 | 0 | 19 | 28 |
| Cebpb | 5 | 8 | 17 | 30 |
| Rxra | 7 | 7 | 16 | 30 |
| Nr1h2 | 19 | 6 | 5 | 30 |
| Ppara | 8 | 14 | 10 | 32 |
| Srebf1 | 13 | 11 | 8 | 32 |
| Hivep2 | 14 | 12 | 9 | 35 |
| Ppargc1a | 16 | 10 | 11 | 37 |
| Bhlhe40 | 17 | 13 | 7 | 37 |
| Ebf1 | 15 | 15 | 12 | 42 |
| Pparg | 11 | 19 | 13 | 43 |
| Mafg | 12 | 18 | 18 | 48 |
| Rxrg | 18 | 16 | 15 | 49 |

**Extended Data Table 3:**
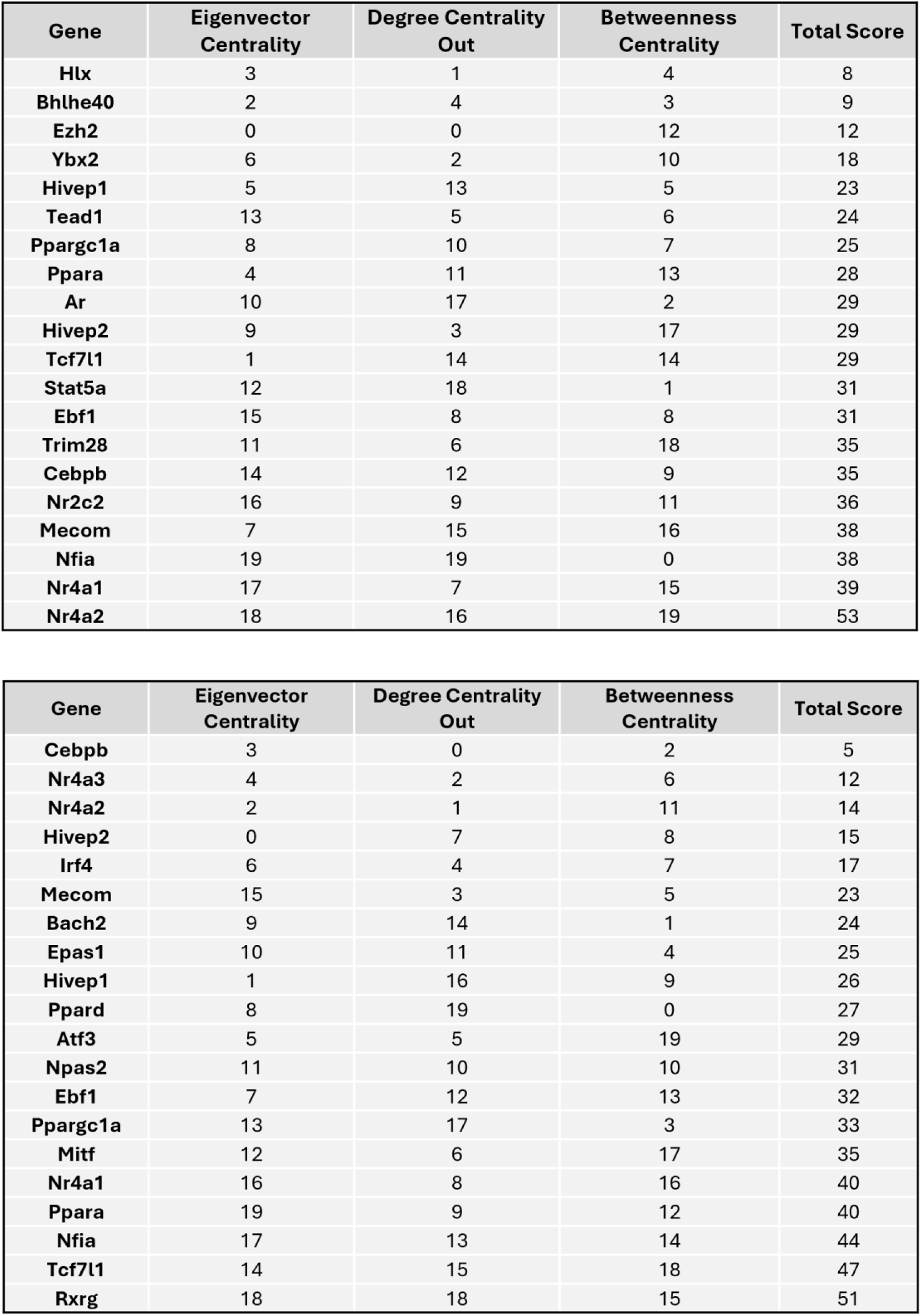
Top transcription factors identified by CellOracle for Males (8W top, 24W bottom)

**Extended Data Figure 1:**
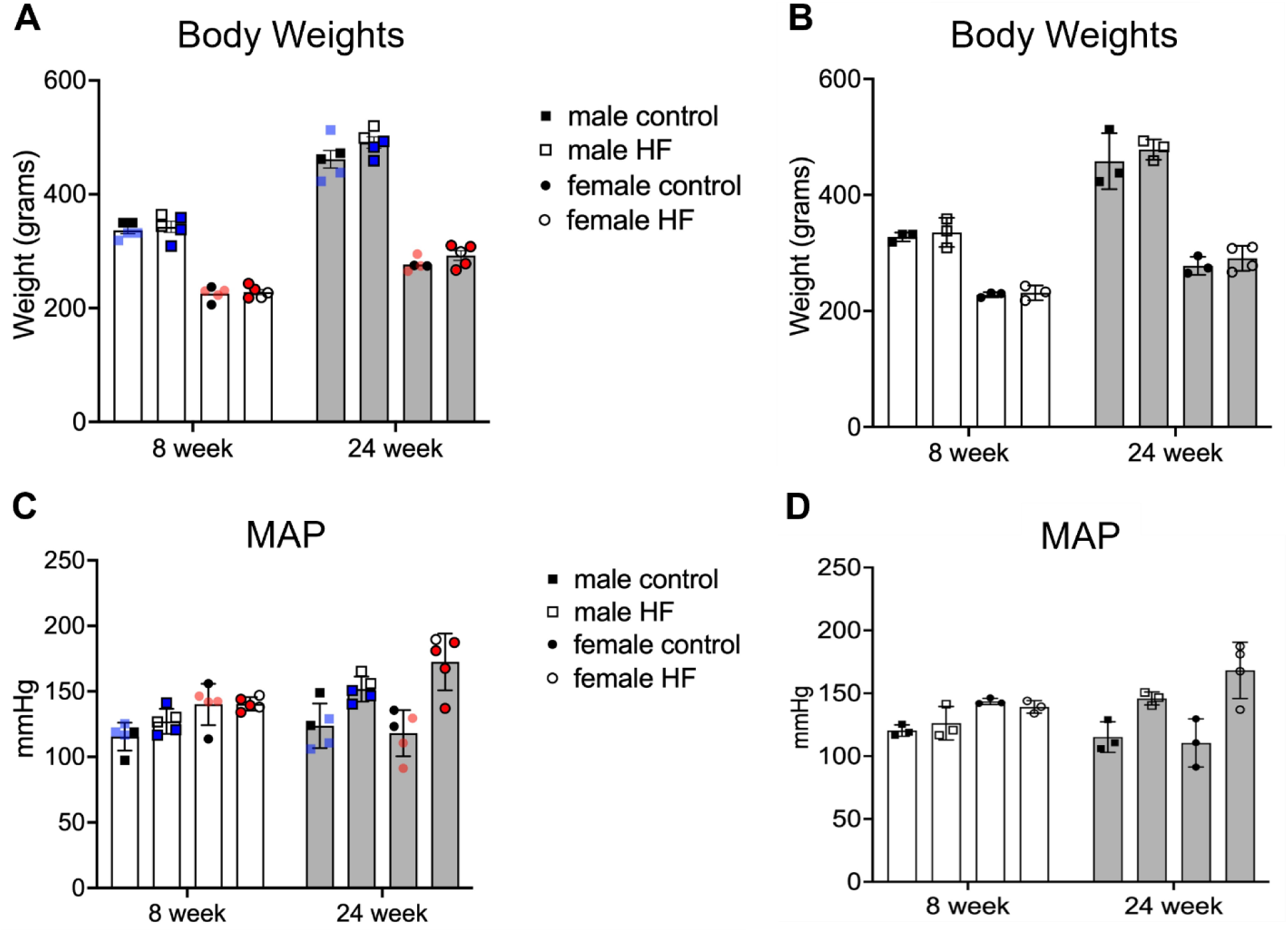
**(A)** Body weights of male and female Dahl SS rats on control and high fat diet in cohorts at 8 (N=5) and 24 weeks (N=5). **(B)** Subset of body weights from rats on control and high fat diet used in reported snRNA-seq experiments at 8 (N=3) and 24 weeks (N=3-4). **(C)** MAP of male and female Dahl SS rats on control and high fat diet in cohorts at 8 (N=5) and 24 weeks (N=5). **(D)** Subset of MAP from rats on control and high fat diet used in reported snRNA-seq experiments at 8 (N=3) and 24 weeks (N=3-4). Colored symbols indicate samples (N=3-4) used for snRNA-seq work. Bars represent means ± SEM.

**Extended Data Figure 2:**
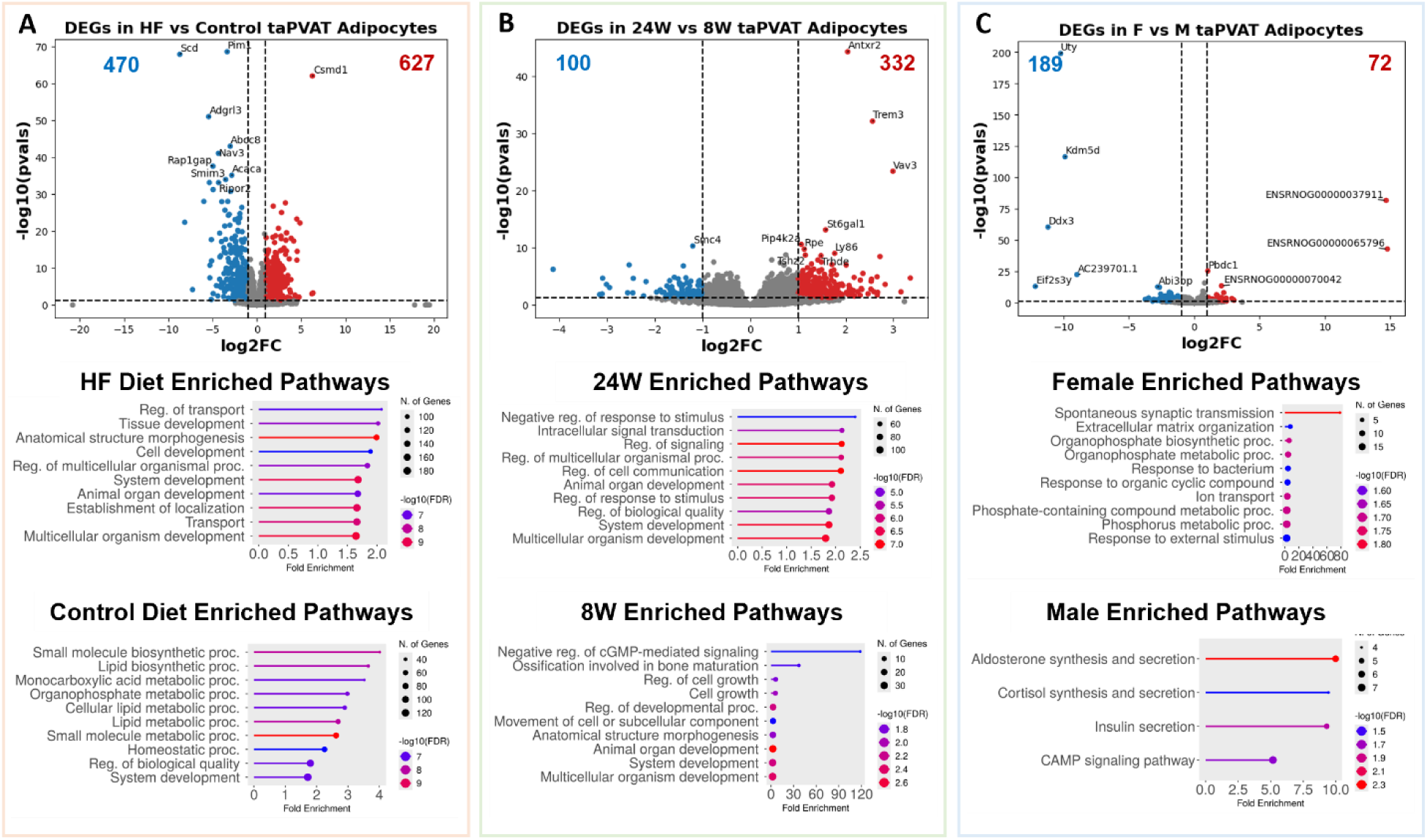
**(A)** (Top) Volcano plot of differentially expressed genes (DEGs) in adipocytes between high fat and control diets. (Middle, Bottom) Enriched GO biological process pathways in rats fed a HF or control diet (includes 8-week, 24-week, M and F data). **(B)** (Top) Volcano plot of DEGs between 8W and 24W diet durations. (Middle, Bottom) Enriched GO biological process pathways in rats on diet for 24-weeks or 8-weeks (includes control diet, HF diet, M and F data). **(C)** (Top) Volcano plot of DEGs between female and male rats. (Middle, Bottom) Enriched GO biological process pathways in female or male rats (includes control diet, HF diet, 8-week, and 24-week data).

**Extended Data Figure 3:**
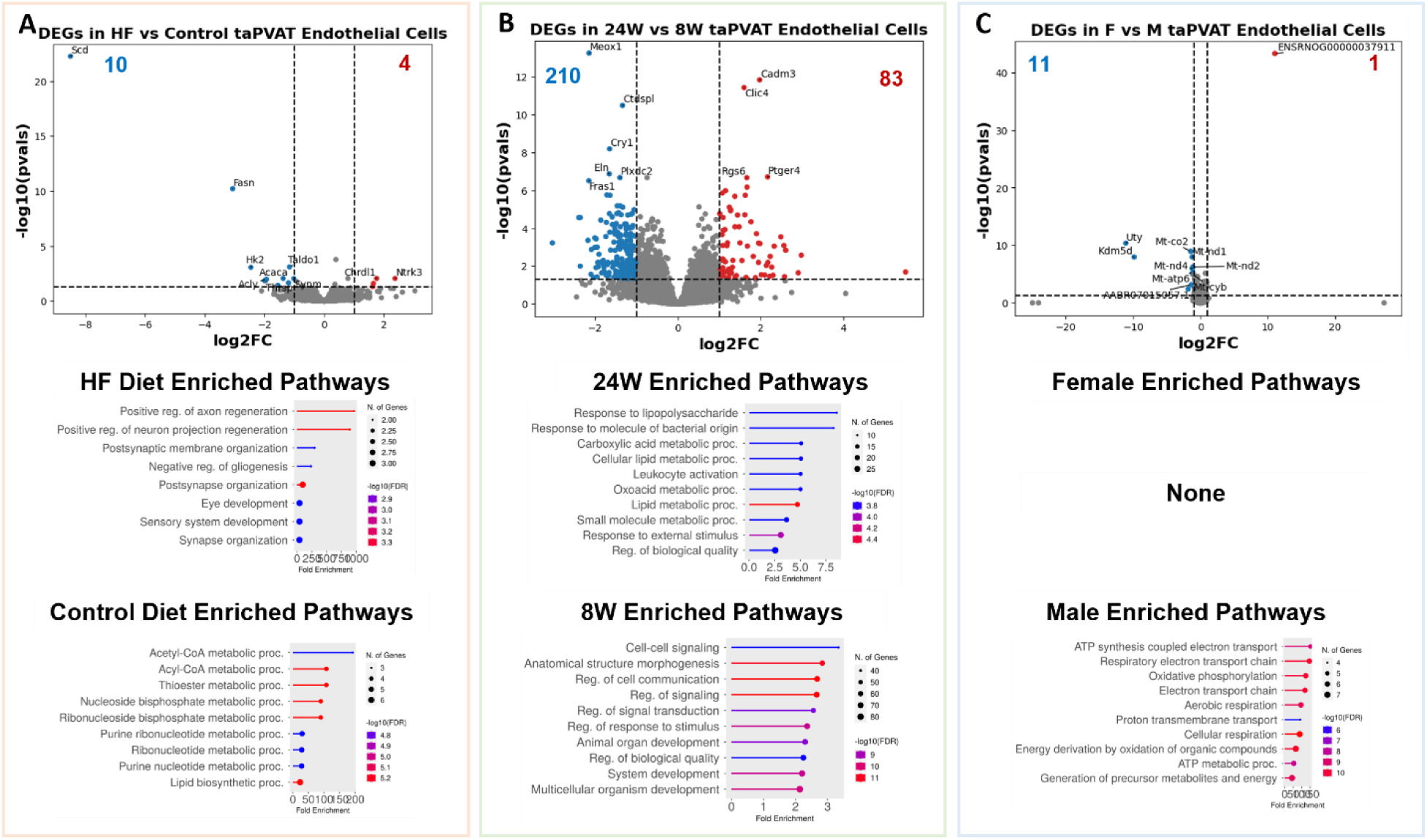
**(A)** (Top) Volcano plot of differentially expressed genes (DEGs) in endothelial cells between high fat and control diets. (Middle, Bottom) Enriched GO biological process pathways in rats fed a HF or control diet (includes 8-week, 24-week, M and F data). **(B)** (Top) Volcano plot of DEGs between 8W and 24W diet durations. (Middle, Bottom) Enriched GO biological process pathways in rats on diet for 24-weeks or 8-weeks (includes control diet, HF diet, M and F data). **(C)** (Top) Volcano plot of DEGs between female and male rats. (Middle, Bottom) Enriched GO biological process pathways in female or male rats (includes control diet, HF diet, 8-week, and 24-week data).

**Extended Data Figure 4:**
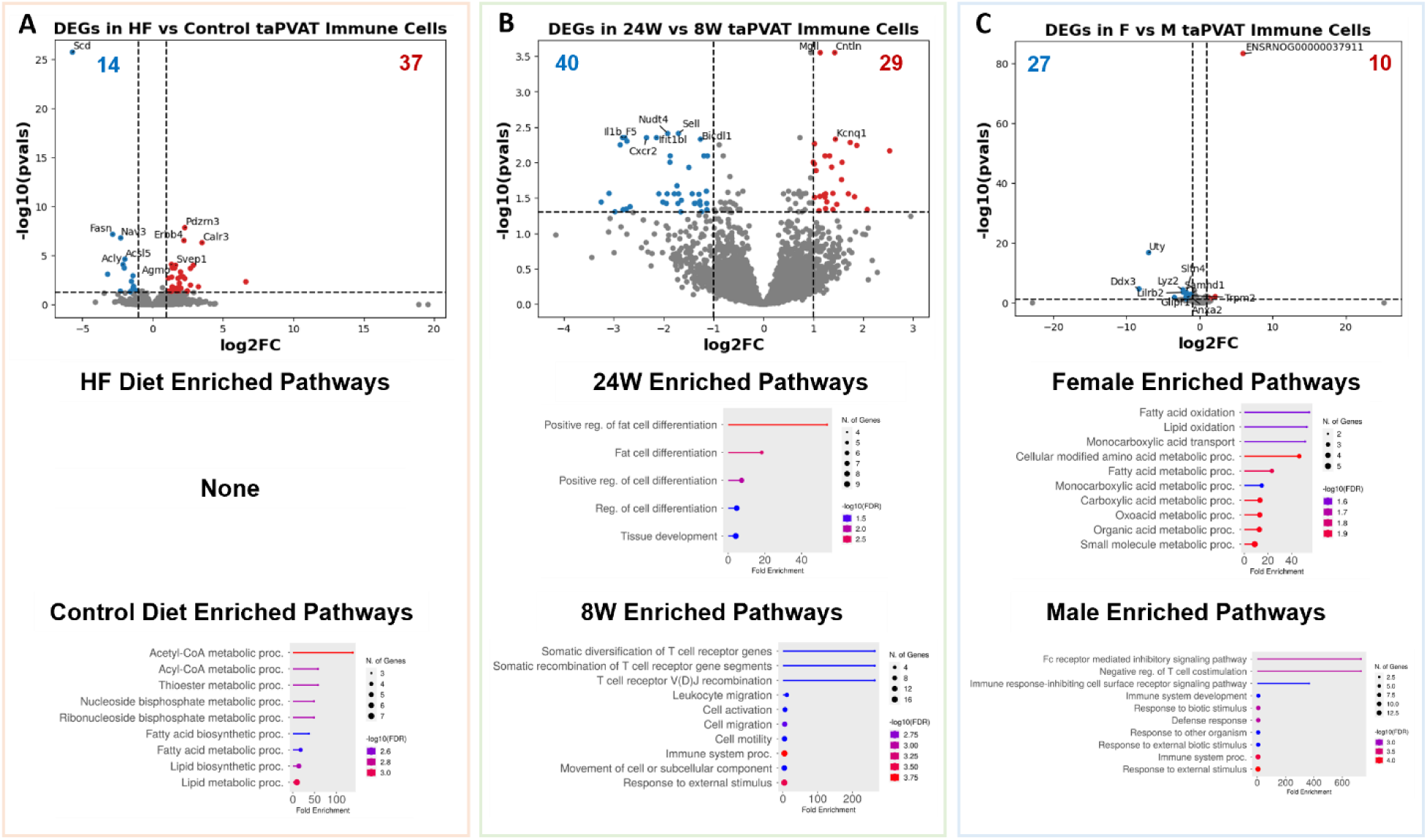
**(A)** (Top) Volcano plot of differentially expressed genes (DEGs) in immune cells between high fat and control diets. (Middle, Bottom) Enriched GO biological process pathways in rats fed a HF or control diet (includes 8-week, 24-week, M and F data). **(B)** (Top) Volcano plot of DEGs between 8W and 24W diet durations. (Middle, Bottom) Enriched GO biological process pathways in rats on diet for 24-weeks or 8-weeks (includes control diet, HF diet, M and F data). **(C)** (Top) Volcano plot of DEGs between female and male rats. (Middle, Bottom) Enriched GO biological process pathways in female or male rats (includes control diet, HF diet, 8-week, and 24-week data).

**Extended Data Figure 5:**
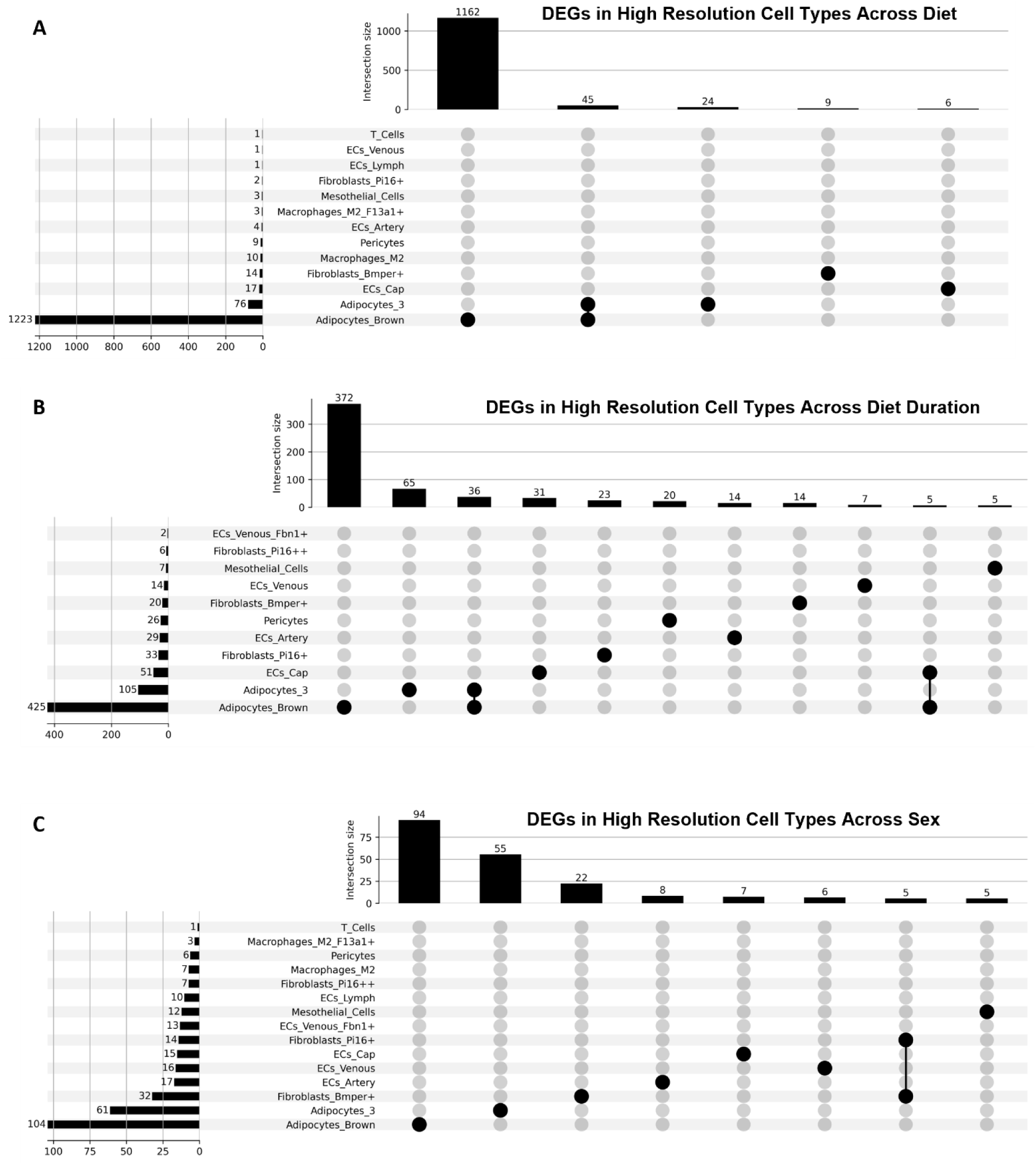
Upset plots of DEGs in low-level cell types across **(A)** diet, **(B)** diet duration, and **(C)** sex.

**Extended Data Figure 6:**
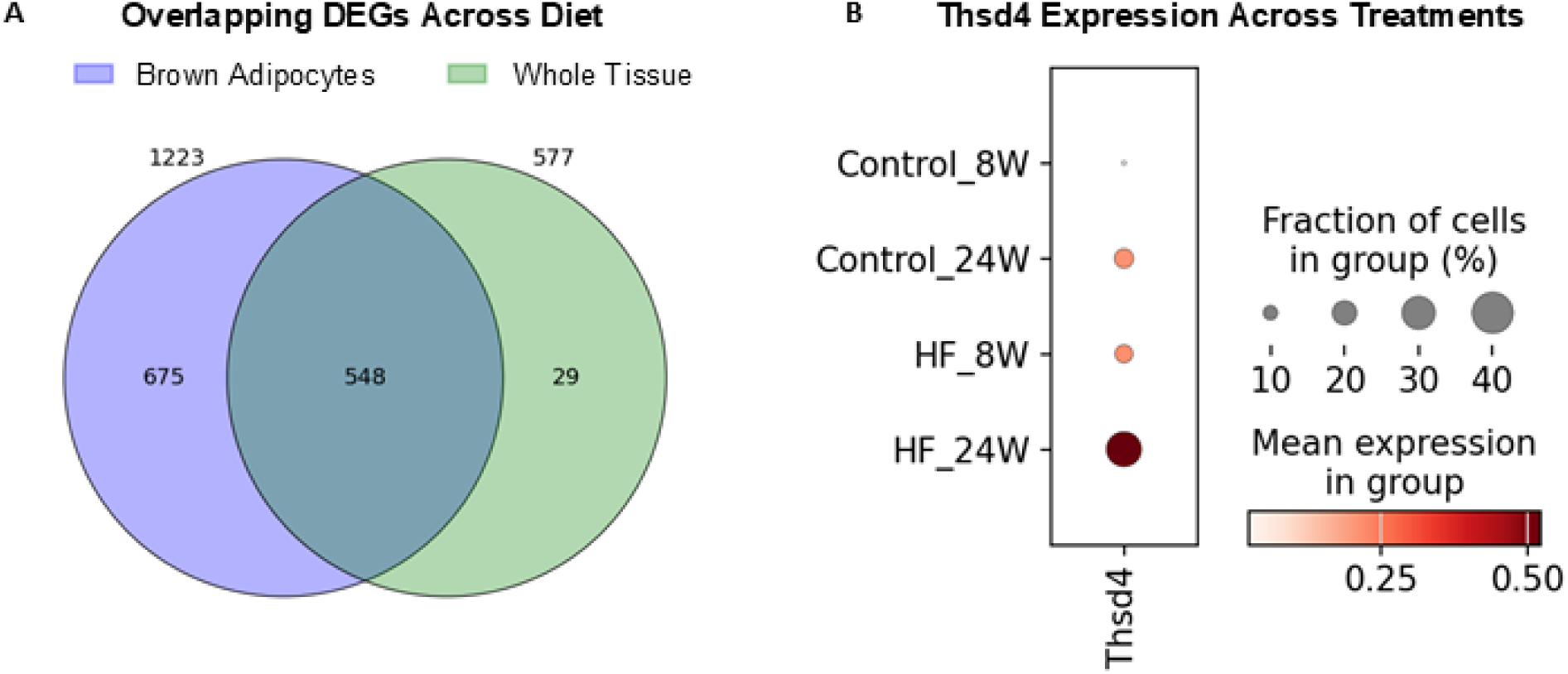
**(A)** Venn diagram highlighting overlapping DEGs between brown adipocytes and the whole tissue when comparing across diets. **(B)** Dot plot of *Thsd4* gene expression across diets and time points. Expression is much higher in the HF diet fed rats and increases in both diets across time from 8W to 24W.

**Extended Data Figure 7:**
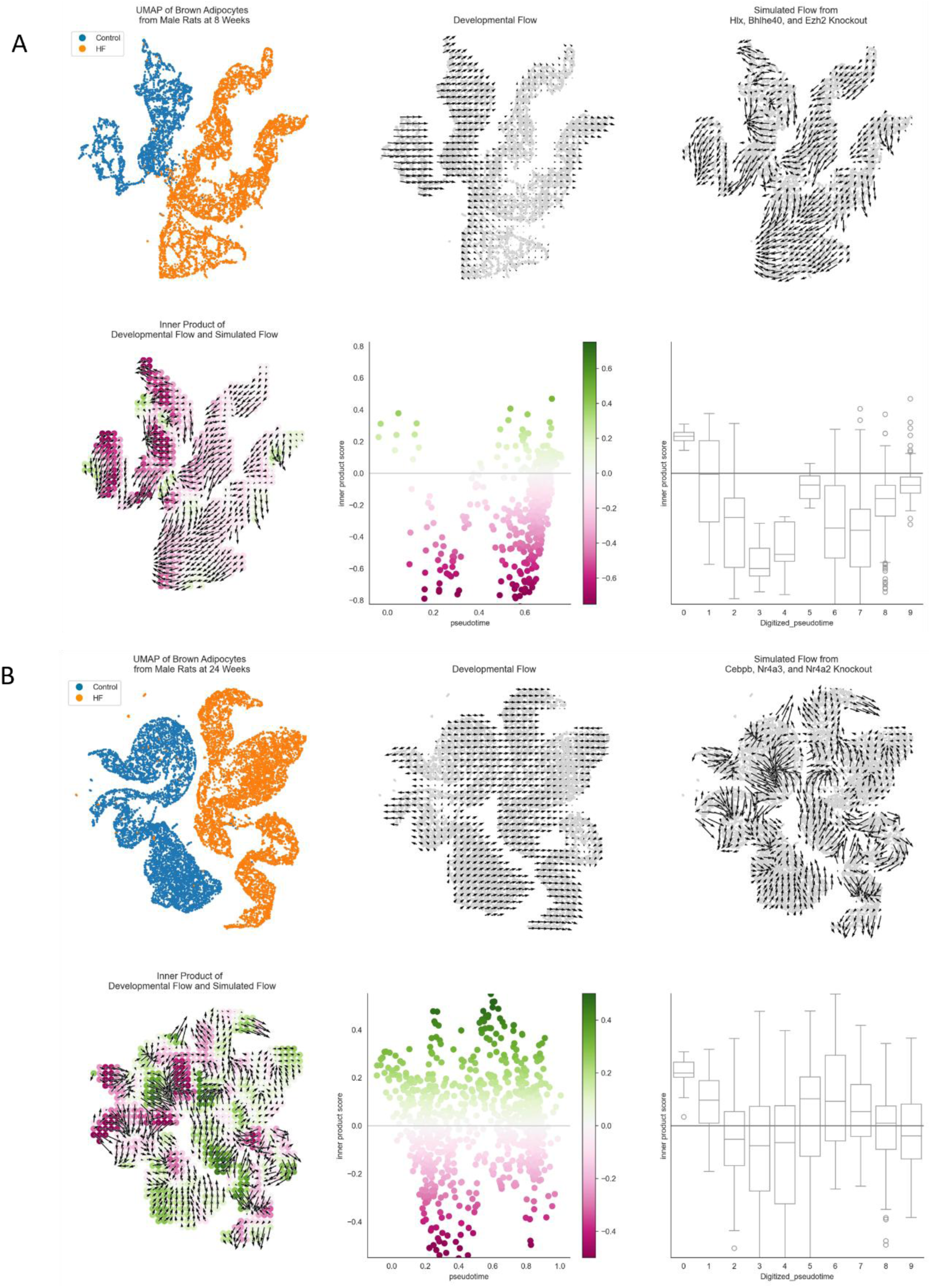
Transcription factor perturbation analyses with CellOracle for **(A)** 8-week males and **(B)** 24-week males.

